# Computational capability of ecological dynamics

**DOI:** 10.1101/2021.09.15.460556

**Authors:** Masayuki Ushio, Kazufumi Watanabe, Yasuhiro Fukuda, Yuji Tokudome, Kohei Nakajima

## Abstract

Ecological dynamics is driven by an ecological network consisting of complex interactions. Information processing capability of artificial networks has been exploited as a computational resource, yet whether an ecological network possesses a computational capability and how we can exploit it remain unclear. Here, we show that ecological dynamics can be exploited as a computational resource. We call this approach “Ecological Reservoir Computing” (ERC) and developed two types of ERC. *In silico* ERC reconstructs ecological dynamics from empirical time series and uses simulated system responses as reservoir states, which predicts near future of chaotic dynamics and emulates nonlinear dynamics. The real-time ERC uses population dynamics of a unicellular organism, *Tetrahymena thermophila*. Medium temperature is an input signal and changes in population abundance are reservoir states. Intriguingly, the real-time ERC has necessary conditions for reservoir computing and is able to make near future predictions of model and empirical time series.

## Introduction

Ecological dynamics are driven by complex interactions. Empirical and theoretical studies have shown that prey-predator, mutualistic, competitive and biotic-abiotic interactions are prevalent, and that they play a vital role in ecological community dynamics (Mougi & Kondoh 2012; Reynolds & Bruno 2013; Ushio *et al*. 2018; Ratzke *et al*. 2020). In nature, the interactions shape an ecological network that includes numerous species (nodes). Information of a node, for example, species abundance, can be processed through interactions and transferred to another node in a very complex way that is often difficult to accurately represent by equations. Population dynamics or community dynamics is temporal fluctuations in species abundance, and is a consequence of the “information processing.” Ecologists have tried to discern rules that govern the ecological dynamics or the information processing.

In computational science, information processing capability of artificial networks is exploited as a computational resource (e.g., artificial neural networks). Artificial neural networks are represented by a network of neuron-like processing units (nodes) interconnected via synapse-like weighted links (interactions), which are typically classified into feedforward neural networks (Schmidhuber 2015) and recurrent neural networks (RNNs) (Mandic & Chambers 2001). A machine learning approach called reservoir computing (RC) is a special type of RNN that is suitable for temporal information processing such as time series analysis (Jaeger 2002; Nakajima & Fischer 2021). In RC, input data are nonlinearly transformed into spatiotemporal patterns in a high-dimensional space by an RNN called a “reservoir.” Then, a pattern analysis from the spatiotemporal patterns is performed in the readout. The main characteristic of RC is that the input weights (***W***_*in*_) and the weights of the recurrent connections within the reservoir (***W***) are not trained, whereas only the readout weights (***W***_*out*_) are trained with a simple learning algorithm such as a linear regression. This simple and fast training process makes it possible to drastically reduce the computational cost of learning compared with standard RNNs, which is the major advantage of RC (Jaeger 2002; Nakajima & Fischer 2021).

Recently, RC implementation using a physical material has been gaining growing attention in machine learning and engineering fields (physical reservoir computing, Nakajima 2020). A nonlinear, complex information processing capability is embedded in a physical material (embodiment, see Pfeifer *et al*. 2007), and thus one can replace a reservoir in RC with a physical material. For example, a soft robotic, octopus-like arm can process an input signal from a motor (i.e., a motor that initiates a movement of the robotic arm), and then the signal transmits through the arm in a way that depends on physical characteristics of the robotic arm such as length, material, and shape. Nakajima *et al*. have shown that such a soft robotic arm has a short-term memory and can be used to solve several computational tasks in real time (Nakajima *et al*. 2014; Nakajima *et al*. 2015).

Several successful examples of physical reservoir computing (Tanaka *et al*. 2019; Nakajima 2020; Nakajima & Fischer 2021) imply that we may be able to exploit information processing capability of other types of networks as a computational resource. Here, we show that ecological dynamics, outputs of ecological networks, can be exploited as a computational resource. We call this approach “Ecological Reservoir Computing (ERC)” and implement two types of ERC in this study (Fig. 1A). The first type of ERC is *in silico* ERC; it reconstructs ecological dynamics from empirical time series using a time-delay embedding (Takens 1981) and simulates the system dynamics in response to input signals. *In silico* ERC uses the reconstructed dynamics and the simulated responses as a reservoir and reservoir states, respectively. Information processing capabilities of reconstructed dynamics were evaluated for prokaryote and fish time series, and this approach successfully predicts the near future of chaotic dynamics and emulates nonlinear dynamics. The results suggest that a real ecological system might also be used as a computational resource. The second type of ERC is real-time ERC; in the present study, we set up an experimental system that enables continuous monitoring of population dynamics of a unicellular eukaryotic organism, *Tetrahymena thermophila*. Medium temperature is used as input and we manipulated the temperature in a small aluminum chamber. Then, we monitored changes in population abundance as reservoir states. Population abundance is an output of this single species system as a result of complex interactions among individuals and environment. Surprisingly, the real-time ERC has necessarily conditions for RC, e.g., echo state property and short-term memory, and is able to make near future predictions of model and empirical time series.

**Figure 1.**
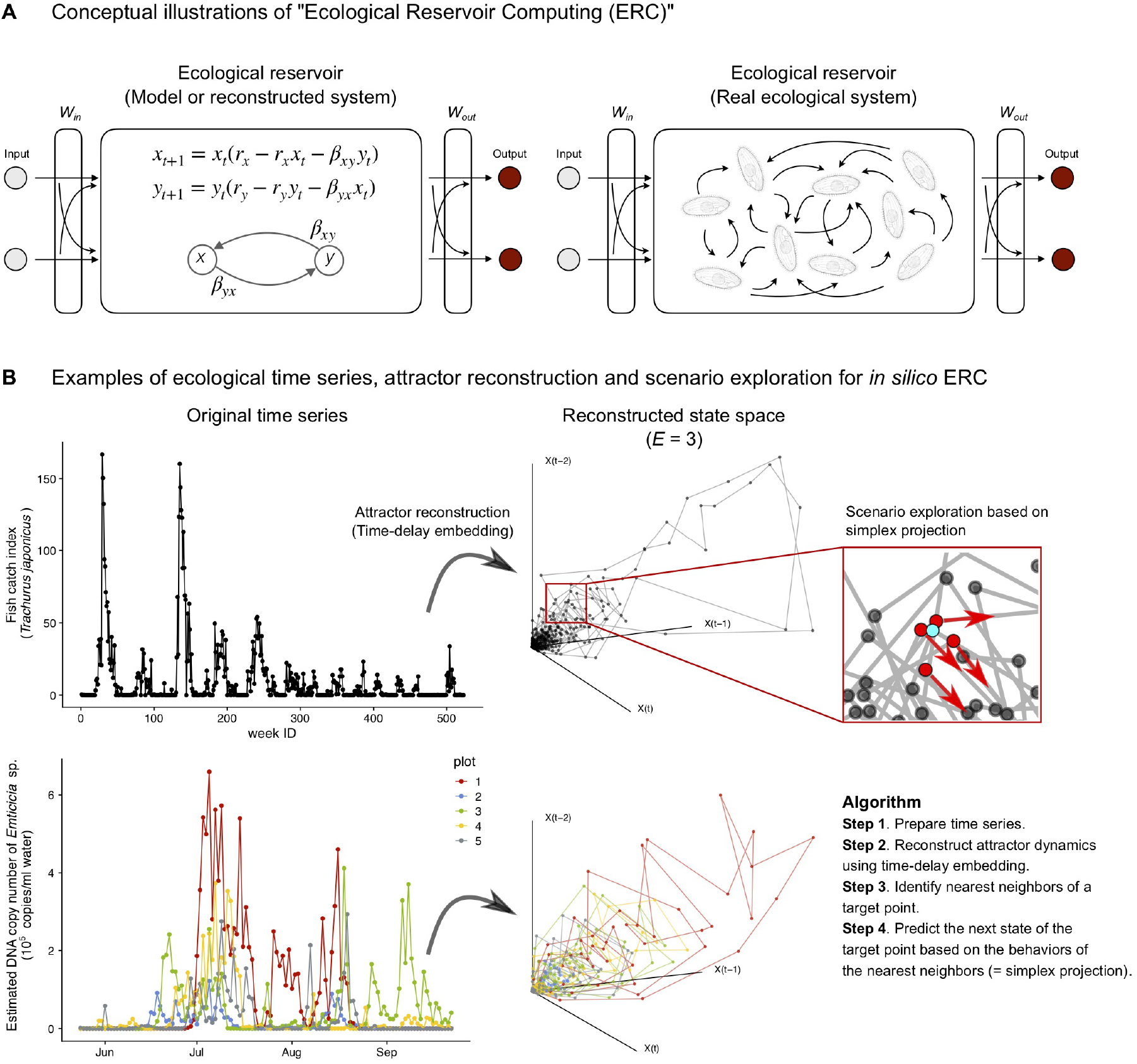
Conceptual illustrations of ecological reservoir computing (ERC) and how *in silico* ERC is implemented. (**A**) Conceptual illustrations of *in silico* ERC and real-time ERC. *In silico* ERC uses either equations or empirical dynamics reconstructed by empirical time series as a reservoir. Real-time ERC uses an empirical ecological interaction network as a reservoir. A node in an ecological reservoir may represent an individual, species, or abiotic variable in this study. (**B**) Examples of ecological time series, state space reconstruction, and scenario exploration for *in silico* ERC. Two empirical time series are shown as examples: Fish catch time series of Japanese jack mackerel (*Trachurus japonicus*) and DNA copy number time series of *Emticicia* sp. in water samples collected from experimental rice plots (Ushio 2020). Empirical attractor dynamics can be reconstructed by time-delay embedding (Embedding dimension = 3). The red inlet indicates that the behavior of a target point (light blue) is predicted by the behaviors (red arrows) of nearest neighbors (red points).

### Ecological reservoir computing: Demonstration of the concept by a toy model

We first demonstrate the concept of ERC using a toy model that is frequently used in ecology. Eqn. [1] shows two coupled difference equations that can be interpreted as a model of two-species population dynamics:

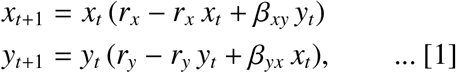

where *x*_*t*_ and *y*_*t*_ indicate a population density of species *x* and *y* at discrete time *t*, respectively. *r*_*i*_ indicates the population growth rate of species *i*, and *β*_*i j*_ indicates influences from species *j* to species *i* (i.e., interspecific interactions). If *β*_*xy*_ and *β*_*yx*_ are negative, the equations represent a two-species competition model (see Fig. S1A for an example of the population dynamics).

The simple nonlinear model can be used as a small reservoir in the context of RC. First, any inputs can be converted using the weight matrix for the input-reservoir connections, ***W***_*in*_, and then reservoir dynamics follow Eqn. [1]. This information processing can be described as follows:

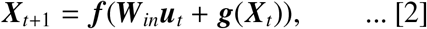

where *t* denotes discrete time, ***X***_*t*_ is the state vector of the reservoir (***X***_*t*_ = {*x*_*t*_, *y*_*t*_}), ***u***_*t*_ is the input vector, and ***g*** is the population dynamics model (***g*** is adjusted so that ***g*** does not show chaotic behavior; Fig. S1A). ***f*** represents an element-wise activation function. While hyperbolic tangent is often used as ***f***, we choose an identity function as ***f*** so that Eqn. [2] can be interpreted as the population dynamics of two species, *x* and *y*, in response to the addition or removal individuals of species *x* and *y*. Then, the reservoir states, ***X***_*t*_, are used to train readout weights, ***W***_*out*_, by a ridge regression (see Methods for the definition of ***W***_*out*_).

This framework enables transforming a traditional ecological population model into an RC system. We used Lorenz attractor as an input (***u***_*t*_; Fig. S1B), and near future predictions of the chaotic time series are indeed possible (Fig. S1C–E). The performance of the small ecological reservoir is low, which is expected given the small reservoir size (the number of reservoir nodes, *N*, is two). However, this example demonstrates how we can potentially exploit ecological dynamics as a computational resource.

### Exploiting reconstructed ecological dynamics as a reservoir

The example of the two-species population model relies on equations, that is, we cannot use the system for RC *in silico* unless equations governing the ecological dynamics are known. Unfortunately, we usually do not know equations that govern real ecological dynamics, and thus, we need an equation-free framework if we want to use ecological dynamics for RC *in silico*.

Scenario exploration, a method to simulate the response of an empirical ecological system to external forces (Deyle *et al*. 2013), may be a promising approach to use empirical ecological time series as a reservoir. Even when equations governing system dynamics are unknown, multivariate system dynamics may be reconstructed from a single time series using a time-delay embedding (Fig. 1B, Takens 1981; Deyle & Sugihara 2011), which is known as state space reconstruction. Also, one may add one or more variables (ordinates) in the reconstructed state space, allowing simulations of ecosystem response to external forces. For example, Deyle *et al*. (2013) predicted how changes in sea surface temperature influence population abundance of Pacific sardine. In the context of RC, changes in sea surface temperature and predicted population abundance of Pacific sardine may be regarded as “input” and “readout,” respectively. The reconstructed state space contains a rule that governs the empirical ecological dynamics and can be exploited as a reservoir.

We demonstrate this method, *in silico* ERC, using empirical ecological time series: (i) fish catch time series collected from pelagic regions in Japan (Fig. 1B, Methods) and (ii) DNA-based quantitative prokaryote time series taken from experimental rice plots (Fig. 1B, Ushio 2020). State spaces were first reconstructed using an optimal embedding dimension determined following a previous study (simplex projection, Sugihara & May 1990). The optimal embedding dimension (*E*) was estimated to be three for both time series, so the state spaces were reconstructed in a three-dimensional space (Fig. 1B). Then, reservoir states were calculated as follows:

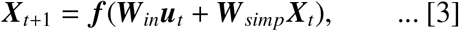

where ***W***_*simp*_ indicates a simplex projection (Sugihara & May 1990), a nonlinear forecasting method that predicts a behavior of a target state based on behaviors of nearest neighbors in the reconstructed state space. The behavior of ***X***_*t*_ is predicted by ***W***_*simp*_ so that ***X***_*t*_ follows the rule of the empirical ecological dynamics (i.e., nonlinear map; see Methods). Then, a hypothetical input, ***u***_*t*_, is added to the state after transformation by ***W***_*in*_ (Fig. S2A). In this case, we again choose an identity function as ***f***. Alternatively, one may apply ***W***_*simp*_ after adding a hypothetical input, ***u***_*t*_, to ***X***_*t*_ as follows:

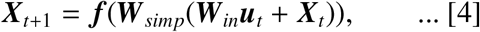

These methods may perform differently depending on the task (In the present study, Eqn. [3] was used for a prediction task, and Eqn. [4] was used for an emulation task and a generation of an autonomous system). In addition, this framework enables multiplexing reservoir states generated by different species (species multiplexing; Fig. S2A). The reconstructed dynamics of different species may generate different reservoir states, which can easily be combined and used to improve the performance of *in silico* ERC.

The *in silico* ecological reservoir possesses a specific memory capacity and shows echo state property (ESP), which are necessary for RC (Fig. S2B–D). We also measured information processing capacities, which can evaluate the expressiveness of the reservoir in terms of memory capacity and nonlinear processing of inputs, using both species systematically (Fig. S3 and Supplementary Text). Then, we tested the performance of *in silico* ERC by several standard tasks: Prediction of chaotic dynamics, emulation of nonlinear autoregression moving average (NARMA) time series, and generation of an autonomous system (Mackey-Glass equation). First, *in silico* ERC with species multiplexing accurately predicts Lorenz attractor (Figs. 2A and S2E). In the *in silico* ERC, DNA-based time series of 500 prokaryotic species that were collected from the same experimental system were multiplexed (Ushio 2020). Interestingly, the prediction accuracy measured by a correlation coefficient increases with the number of species multiplexed (Fig. 2B), suggesting that species diversity of a community might be related to the computational capability of an ecological community. Second, NARMA2 can be accurately emulated with species-multiplexed *in silico* ERC (Fig. 2C), although the performance is lower than that of a typical RC, echo state network (ESN) (for details of the parameter setting of ESN, see Supplementary Text). Third, the Mackey-Glass equation cannot be embedded in a closed-loop with species-multiplexed *in silico* ERC in our current numerical experiments, but in silico ERC generates different attractor dynamics (Fig. 2E, F).

**Figure 2.**
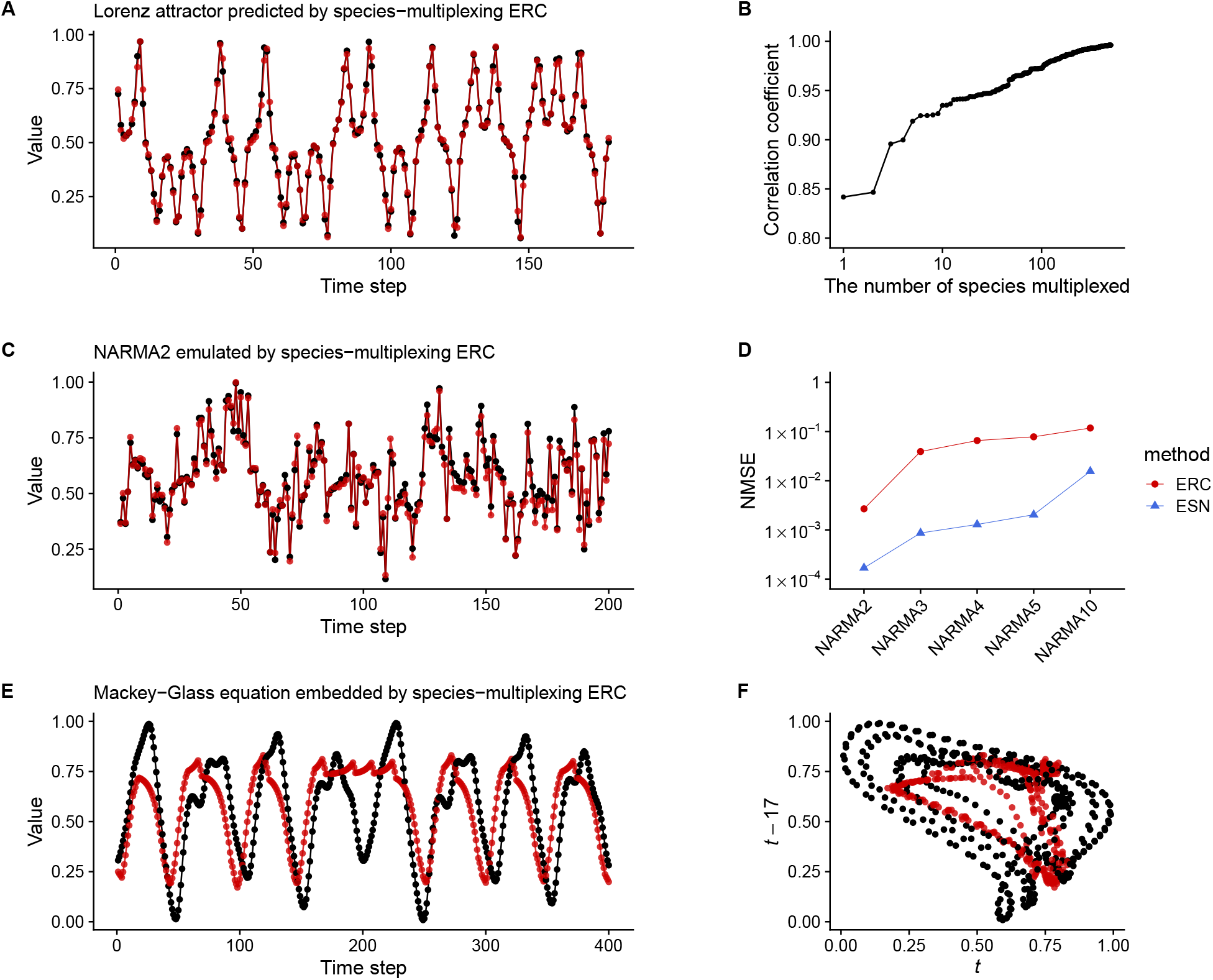
The performance of species-multiplexed *in silico* ecological reservoir computing (ERC). (**A**) Time series of Lorenz attractor (black points and lines) and one time-step future predictions by species-multiplexed *in silico* ERC (red points and lines). (**B**) Correlation coefficients of observed and predicted values of Lorenz system and the number of species multiplexed. (**C**) Nonlinear autoregression moving average (NARMA) time series (black points and lines) and emulation by species-multiplexed *in silico* ERC (red points and lines). NARMA is NARMA2 in this panel (**D**) Normalized mean square error (NMSE) of the NARMA emulations by species-multiplexed *in silico* ERC (red points and lines) and echo state network (ESN; blue points and lines). (**E**) The closed-loop embedding of the Mackey-Glass equations by species-multiplexed *in silico* ERC. The original attractor (black points and lines) was learned by the *in silico* ERC and autonomous dynamics was generated from time point zero (red points and lines). (**F**) Two-dimensional representation of the original Mackey-Glass attractor (black points) and that generated by the *in silico* ERC (red points).

Interestingly, the analysis on information processing capacity suggests that the *in silico* ERC has relatively higher nonlinear processing capacity than the linear one compared with the profile of the conventional ESNs (Fig. S3C). Altogether, these results show that the method based on scenario exploration can be used as RC and solve several standard tasks. More importantly, *in silico* ERC may be useful to screen for real-time ecological dynamics with a high computational capability, and the successful applications of *in silico* ERC suggest that real-time ecological dynamics may also be exploited as a computational resource.

### Real-time ecological reservoir computing

*In silico* ERC suggests that we may exploit real-time ecological dynamics as a computation resource. Here we show that RC is possible even with real ecological dynamics. We focused on the microbial dynamics in a culture system because the system can be easily intervened in and monitored. A eukaryotic unicellular organism, *Tetrahymena thermophila* (hereafter, *Tetrahymena*), is a model organism for molecular biology (Cassidy-Hanley 2012). It usually grows on a bacterial diet, but can easily be cultured on inanimate media (Cassidy-Hanley 2012). Also, it has a high growth rate (the doubling time under optimal conditions is *ca*. two hours), and its growth rate varies depending on environmental factors such as temperature and nutrient concentrations (Fig. S4–S5).

We set up an experimental system to use *Tetrahymena* population dynamics as a reservoir (Fig. 3A–D). The *Tetrahymena* population is cultivated on modified Neff medium (see Methods) and transferred to an aluminum chamber (Fig. 3B–D) when used for an experiment. Medium temperature is accurately controlled with a custom temperature-regulator (Fig. 3A), which is an input signal of real-time ERC. *Tetrahymena* population dynamics is monitored by taking time-lapse images from the bottom of the incubator, and the number of cells is counted by a standard particle analysis (Fig. S6). Although there is only a single species in the system, many factors, including temperature, medium concentration, cell-to-cell interactions, and individual behaviors affect cell numbers captured at the bottom of the chamber (Fig. 3E). Indeed, previous studies have demonstrated that population dynamics of *Tetrahymena* is governed by complex, nonlinear processes (Becks *et al*. 2005; Jordan *et al*. 2013; Weisse *et al*. 2016). Thus, the number of cells captured in an image is a response of the system to an external force, temperature. Specifically, the cell dynamics can be formulated as:

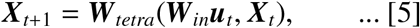

where ***W***_*in*_ determines how the effects of temperature, ***u***_*t*_, propagate to the population dynamics, and ***W***_*tetra*_ determines how temperature influence (***W***_*in*_***u***_*t*_) and population density captured at the bottom of the chamber (***X***_*t*_) interact in the chamber. Importantly, we do not know exact formulations of ***W***_*tetra*_ and ***W***_*in*_, but we can still use this system for RC if ***W***_*tetra*_ and ***W***_*in*_ are time-invariant (note that ***W***_*tetra*_ and ***W***_*in*_ could be nonlinear maps that represent rules governing empirical dynamics).

**Figure 3.**
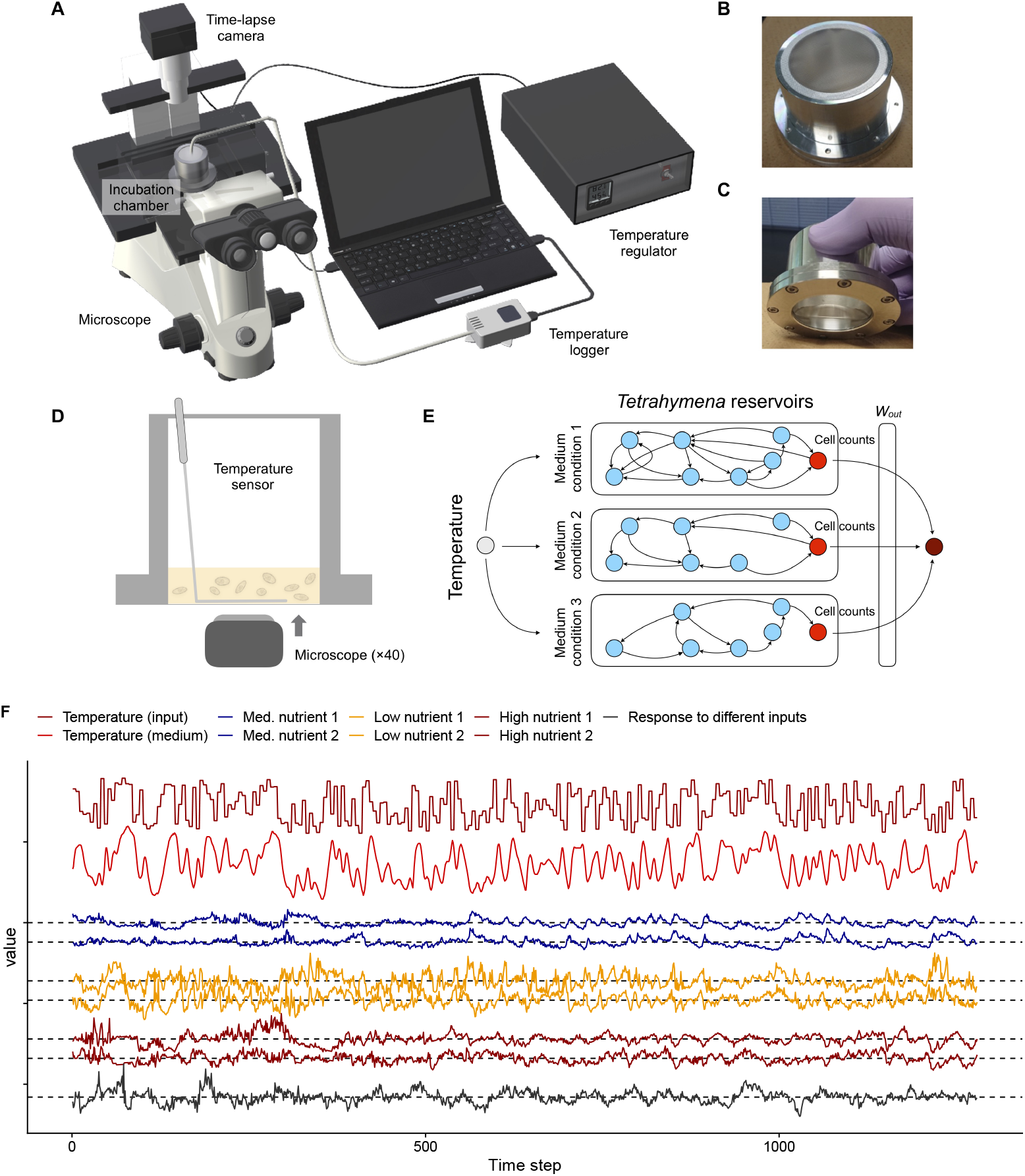
The experimental set up of *Tetrahymena thermophila* reservoir. (**A**) Illustrations of experimental set up. (**B, C**) Pre-incubated *Tetrahymena* population is maintained in an aluminum chamber. The total medium volume is five ml, and the concentration of nutrient is adjusted to change parameters of the population dynamics. (**D**) Cell images were taken from the bottom of the chamber. The number of cells was counted using a custom image analysis pipeline. (**E**) We used 1.6%, 4%, and 10% modified Neff medium in the experiments. Temperature information first transmits from the regulator to the aluminum chamber, and then propagates to several nodes in the medium and *Tetrahymena*. The temperature information is processed through complex interactions among temperature, medium, and behavior and physiology and *Tetrahymena*. The number of cells at the bottom of the chamber may contain the processed information and we use it as a reservoir state. Reservoir states of different nutrient concentrations were used to improve the performance of real-time ERC (i.e., space multiplexing) (**F**) Time series indicate input temperature (dense red), medium temperature (red), and a population density index (relative residual of the population density) in a 4% (blue line; Low nutrient), 1.6% (orange line; Med. nutrient), or 10% (brown line; High nutrient) modified Neff medium. Black line indicates population density index of *Tetrahymena* in a 4% modified Neff medium in response to a different temperature input sequence.

We first tested whether the *Tetrahymena* reservoir has a memory capacity and echo state property (ESP). Uniform random values were set in the temperature regulator, and medium temperature and *Tetrahymena* populations were monitored. Temperature inputs were switched every five minutes, and each image was taken every minute (Fig. 3F). As the maximum number of inputs is 256 for the experimental system, the monitoring continued for 21 hours 20 minutes, generating 1280 images. To make the time series stationary and unbiased, we first estimated the trend by an additive model (Wood 2004), and calculated the relative residuals from the trend (Fig. S6). The relative residuals were used an index of *Tetrahymena* population density, and used as reservoir states hereafter (For more details, see Methods and Supplementary Text). An example of the monitoring results with the same input sequence for different trials is shown in Figs. 3F and S7 (see also Supplementary Video 1, https://youtu.be/z_QeEka4W3w), which shows a clear common-signal-induced synchronization that is a signature of ESP (Lu *et al*. 2018; Inubushi *et al*. 2021).

For RC, two strategies were adopted to increase the reservoir size: time-multiplexing and spatial-multiplexing using different medium concentrations. Five cell counts (one-minute interval) were acquired for one temperature input (five-minute interval); those five were used as the reservoir state (time-multiplexing). Also, to increase the reservoir size, the uniform random temperatures were inputted to the *Tetrahymena* population in different medium concentrations (i.e., 1.6%, 4%, and 10% modified Neff medium). The growth rate depends on the medium concentrations (Fig. S5), suggesting that ***W***_*tetra*_ is a function of the medium concentration, and therefore, the population dynamics under a different medium concentration may be used as a reservoir with different parameters (Fig. 3E).

We tested three medium concentrations and ran two experiments for each medium concentration (Fig. 3E). The same medium concentration generated similar population dynamics (Fig. 3E). We further tested the correspondences between the two runs for each medium concentration, and found that the state differences (i.e., the absolute distance between two states) become smaller when the same input sequence is inputted to the system (Figs. 3F and S7A–F). On the other hand, with a different input sequence in the 4% medium system, the population dynamics show a different pattern, and the state difference does not converge (Figs. 3F and S7G–H). These results suggest that the system has ESP (Lu *et al*. 2018; Inubushi *et al*. 2021). In addition, the population dynamics have a specific memory capacity; the dynamics recover the input values at 5–15 minutes ago (= 1–3 steps ago) (Fig. 4A–C). These characteristics enable the *Tetrahymena* reservoir to measure the medium temperature (Fig. 4D, E), showing that, by using a short-term memory of the reservoir, the *Tetrahymena* population dynamics can work as a “thermometer” of the system. Together, these results suggest that the ecological reservoir may be exploited as a computational resource.

**Figure 4.**
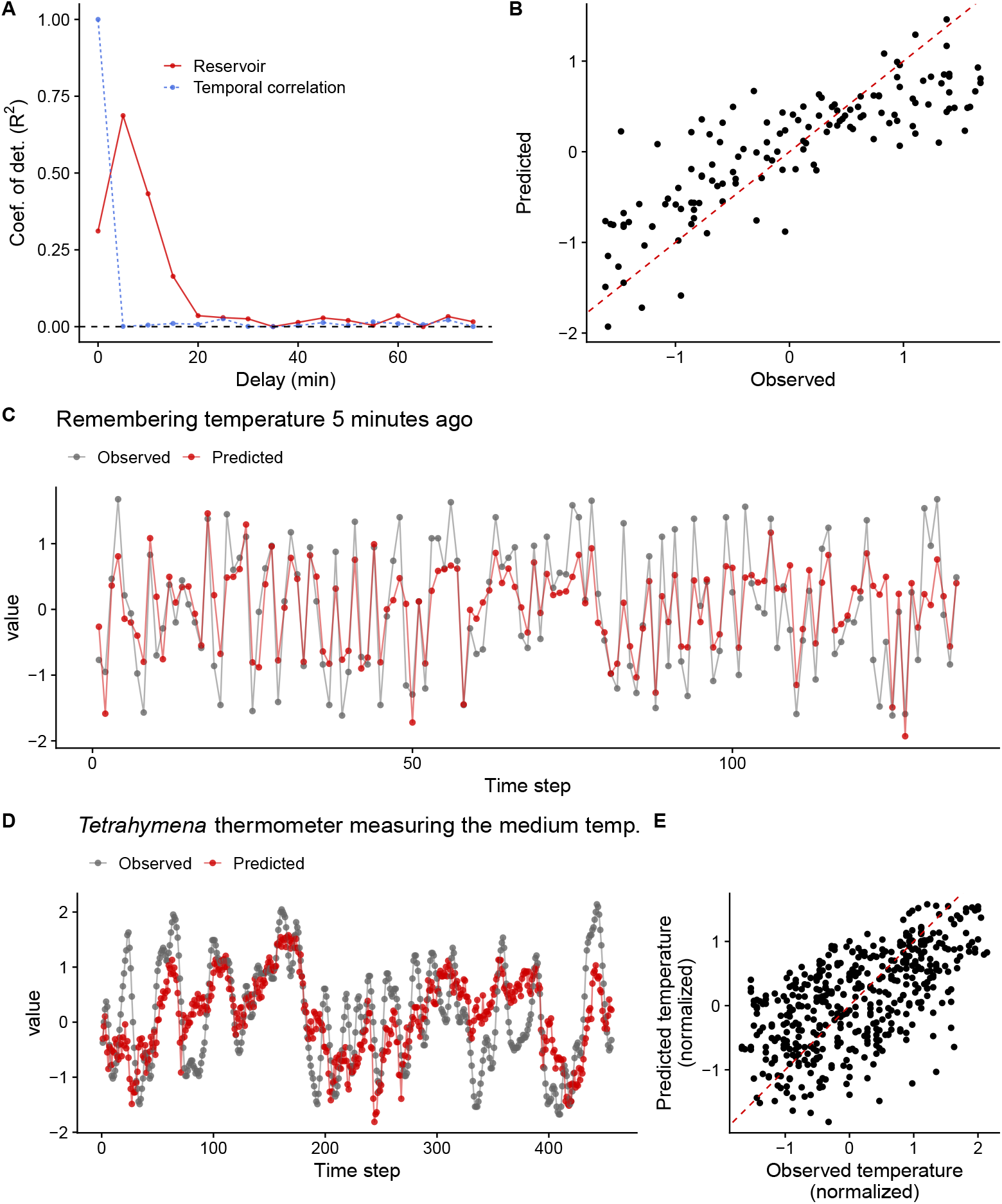
Memory capacity of the *Tetrahymena* reservoirs. (**A**) Memory capacity of the *Tetrahymena* reservoir measured using three time series of population density index (each from three medium concentrations) as a training set and the other three time series as a test set. Red points and lines indicate how well the *Tetrahymena* dynamics remembers the uniform random inputs. Blue points and lines indicate temporal correlations. (**B**) Correlations between observed and predicted values of uniform random inputs. Red dashed line indicates 1:1 line. (**C**) Observed (gray points and lines) and predicted (red points and lines) time series of uniform random inputs. In the panel **C**, temperatures five minutes ago were predicted by the states of the *Tetrahymena* reservoir. (**D**) Measurements of the medium temperatures five minutes ago by the *Tetrahymena* reservoir, suggesting that the *Tetrahymena* reservoir may work as a thermometer (*Tetrahymena* thermometer). (**E**) Correlations between observed and predicted temperature by the *Tetrahymena* thermometer.

To explicitly show that the *Tetrahymena* reservoir can solve computational tasks, we predicted three time series: Lorenz attractor (model time series) and two fish-catch time series (empirical time series; see Methods). As with the uniform random inputs, the same inputs generate similar population dynamics under the same medium concentration, showing ESP of the system (Fig. S8). By time-multiplexing those reservoir states, the *Tetrahymena* reservoir reasonably predicts the near future of the three time series (Figs. 5A–C; see Supplementary Video 2 for how the *Tetrahymena* reservoir predicts the near future, https://youtu.be/SUmkYAnfjFk). The predictions made by the *Tetrahymena* reservoir are more accurate than those made by linear readout (i.e., a ridge regression) at certain time points, suggesting that the computational capability of *Tetrahymena* population dynamics has been successfully exploited by the experimental system. The *Tetrahymena* reservoir predicts 15 time-step future of Lorenz attractor, 19 time-step future of flatfish time series, and 30 time-step future of Japanese jack mackerel time series (Fig. 5D–F; for a version that uses the medium temperature as input, see Fig. S9D–F).

**Figure 5.**
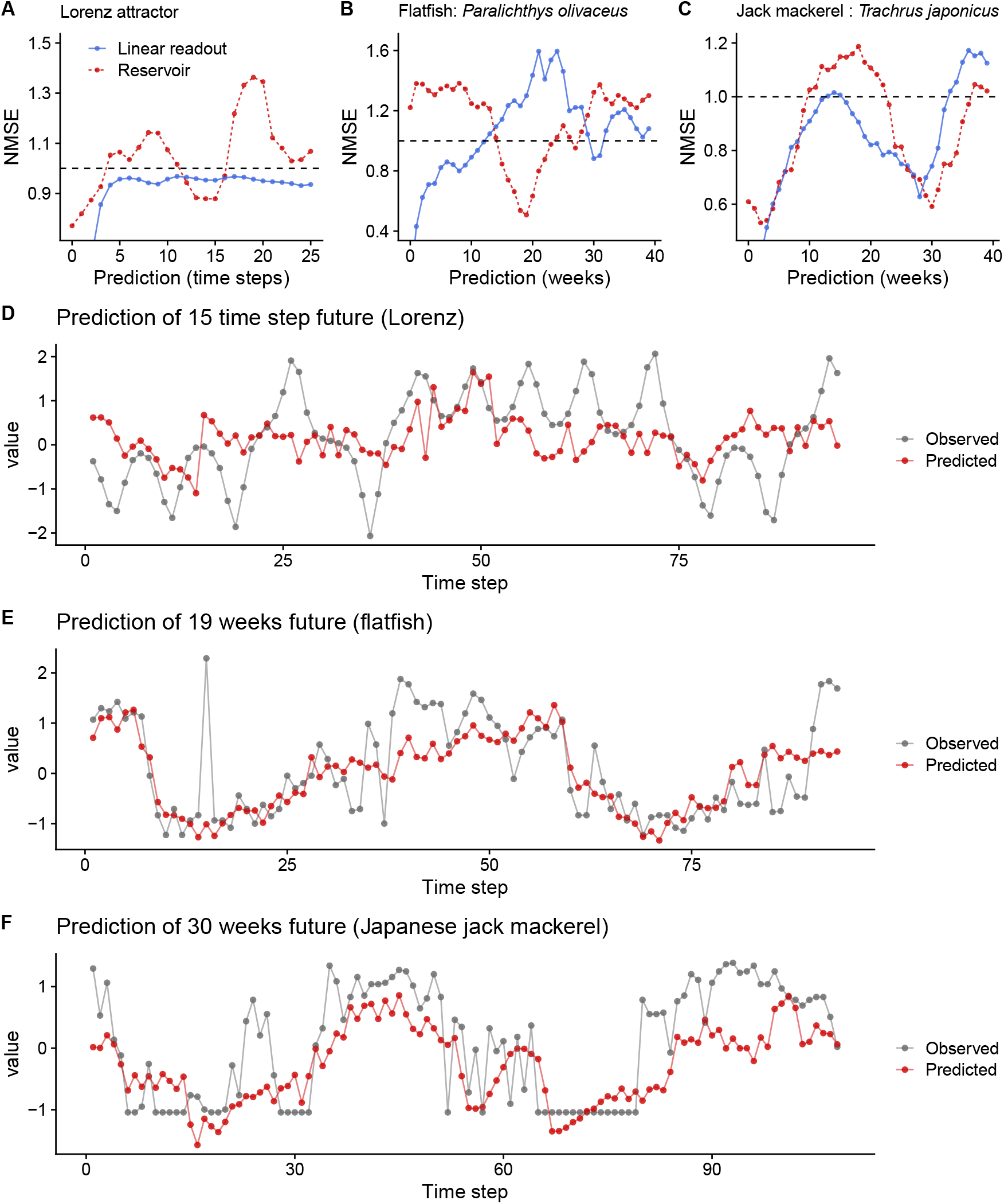
The performance of the *Tetrahymena thermophila* reservoir in predicting model and empirical time series. The relationship between prediction accuracy (Normalized Mean Square Error; NMSE) and prediction time step of (**A**) Lorenz attractor, (**B**) flatfish time series, and (**C**) Japanese jack mackerel time series. Red points and lines indicate predictions by the *Tetrahymena* reservoir, and blue points and lines indicate predictions by ridge regressions. NMSE below 0.4 was not shown in **A**-**C**.Time series of observed (gray points and lines) and predicted (red points and values) of (**D**) Lorenz attractor with 15 time-step future prediction, (**E**) flatfish time series with 19 weeks future prediction, and (**F**) Japanese jack mackerel time series with 30 weeks future prediction.

### Concluding remarks

In the present study, we show a proof-of-concept that ecological dynamics can be exploited as a computational resource in two ways: *in silico* ERC and real-time ERC. The former provides a numerical framework to quantify the potential computational capability of the ecological dynamics and to use the reconstructed dynamics as a computational resource. The framework also provides a unique opportunity to screen for ecological dynamics with a high computational capability, which may be useful for searching for potentially high-performance real ecological dynamics. Furthermore, quantifying the computational capability of existing ecological time series and comparing the computational capabilities with ecological factors (e.g., species identity, phylogeny, and environmental factors) would be an interesting direction. The latter is more intriguing, and its significance is multi-fold. First, real-time ERC is a novel computational framework. Though the computational performance of ERC is still poorer than that of the typical RC, other ecological dynamics (e.g., high-diversity community dynamics) with different experimental settings (e.g., different input signals such as light) will possess different reservoir properties (e.g., with/without echo state property and different memory capacity), and such ecological reservoirs might outperform the typical RC. Second, the real-time ERC enables quantifications of the computational capability of empirical ecological populations or communities. The computational capability may be regarded as an “extended” functional trait of organisms, which should evolve by interacting with biotic and abiotic factors in a natural habitat. Identifying responsible genes for the computational capability would be a fascinating direction. Third, a community with high diversity may potentially have a high reservoir size, which could possess a high computational capability (as shown in Fig. 2B). The potential positive relationship between community diversity and computational capability may add a new value to biodiversity. Lastly, if the “closed-loop” approach as shown in the Mackey-Glass equation in *in silico* ERC (Fig. 2E, F) is successful in real-time ERC, it would imply that we may be able to design specific dynamics in real-time ecological dynamics. It would be a novel approach to manipulate ecological dynamics. Altogether, this study presents how ecological dynamics can be exploited as a computational resource and how ERC provides a new way for understanding, exploiting, and managing ecological dynamics.

## Methods

### A classic reservoir computing framework: Echo state network (ESN)

In the early 2000s, Echo State Networks (ESNs) as well as Liquid State Machines (LSMs) were proposed as a seminal Reservoir Computing (RC) approach (Jaeger 2002; Maass *et al*. 2002). ESNs (and LSMs) are different from conventional Recurrent Neural Networks (RNNs) in that weights on the recurrent connections in the reservoir are not trained but only the weights in the readout are trained (Jaeger 2002). To apply a simple machine learning method to the readout, the reservoir should be appropriately designed in advance. The characteristics of ESNs are briefly described below.

The ESN model was proposed by Jaeger (Jaeger 2002; Jaeger *et al*. 2007). This model uses an RNN-based reservoir consisting of discrete-time artificial neurons. When the feedback from the output to the reservoir is absent, the time evolution of the neuronal states in the reservoir is described as:

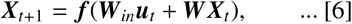

where *t* denotes discrete time, ***X***_*t*_ is the state vector of the reservoir units, ***u***_*t*_ is the input vector, ***W***_*in*_ is the weight matrix for the input-reservoir connections, and ***W*** is the weight matrix for the recurrent connections in the reservoir. The function ***f*** represents an element-wise activation function of the reservoir units, which is typically a sigmoid-type activation function. Eqn. [6] represents a non-autonomous dynamical system forced by the external input ***u***_*t*_. The output is often given by a linear combination of the neuronal states as follows: ***z***_*t*_ = ***W***_*out*_ ***X***_*t*_, where ***z***_*t*_ is the output vector and ***W***_*out*_ is the weight matrix in the readout. In supervised learning, this weight matrix is trained to minimize the difference between the network output and the desired output for a certain time period.

### ERC with a two-species model system: Demonstration of the concept of ERC

To demonstrate the concept of Ecological Reservoir Computing (ERC), we first used logistic equations, a commonly used system to simulate ecological population dynamics, for RC. Eqn. [1] in the main text shows two coupled difference equations that can be interpreted as a model of two-species population dynamics. In the demonstration, we used the following equations (corresponds to Eqn. [1] in the main text):

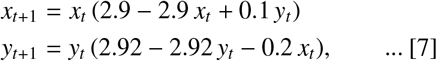

The simple nonlinear model can be used as a small reservoir in the context of RC, as in the main text. Parameter values used in the toy model are follows: *r*_*x*_ = 2.9, *r*_*y*_ = 2.92, *β*_*xy*_ = 0.1, *β*_*xy*_ = −0.2. The sparsity (i.e., the proportion of zero in the matrix elements) of ***W***_*in*_ was set to 0.1. Matrix elements of the input weight were chosen from a uniform random distribution, and they were multiplied by 0.2 to adjust the influence of the input vectors. Readout was trained using a ridge regression (*λ* for the regularization = 0.05).

### In silico *ERC: Empirical ecological time series and scenario exploration*

Empirical ecological time series is used to simulate an ecological system’s responses (for how the ecological time series obtained, see Supplementary Text). For this purpose, we developed a framework based on state space reconstruction (SSR), a method to reconstruct an original dynamics from a single time series (Takens 1981; Deyle & Sugihara 2011) (Fig. 1B). In a natural ecosystem, it is often impossible to collect time series of all potentially important variables involved in a target ecosystem dynamics. Takens (1981) offered a theoretical basis to solve this problem: Takens’ embedding theorem demonstrated that a shadow version of the attractor can be reconstructed using a single observed time. In other words, delineation of the system state trajectory, originally constructed using multiple variables, can be possible even if a time series is available only for a single variable (Takens 1981; Sauer *et al*. 1991). To embed such a single time series, *x*_*t*_, vectors in the putative phase space are formed from time-delayed values of the time series, {*x*_*t*_, *x*_*t*−*τ*_, *x*_*t*−2*τ*_, …, *x*_*t*−(*E*−1)*τ*_}, where *E* is the embedding dimension, and *τ* is the time-lag unit. This procedure, the reconstruction of the original dynamics, is known as SSR.

Based on SSR, Deyle et al. (2013) proposed a numerical method to predict an ecosystem’s responses to external forces (Fig. 1B). A typical way to view temporal dynamics is as separate time series of each species. For example, see Fig. 1B for time series of Japanese jack mackerel (*Trachurus japonicus*) and bacteria (*Emticicia* sp.). However, according to Takens’ theorem, system dynamics may be delineated by plotting time-lagged coordinates in a multidimensional space. In the case of Japanese jack mackerel and *Emticicia* sp., the system dynamics can be reconstructed by taking three coordinates (*E* = 3; the optimal *E* was estimated by simplex projection), ***X***_*t*_ = {*x*_*t*_, *x*_*t*−1_, *x*_*t*−2_}. The reconstructed attractor preserves essential information about the system dynamics, and we may be able to predict near future behaviors of the system (Fig. 1B). One or some of the coordinates may be replaced with an external force (*y*_*t*_) such as temperature, generating a new, but topologically equal, attractor, e.g., ***X***_*t*_ = {*x*_*t*_, *x*_*t*−1_, *y*_*t*_}. A hypothetical input, for example, ***W***_*in*_***u***_*t*_ = {0, 0, *u*_*t*_}, may be added to the vector, and the behavior of the perturbated vector, ***W***_*in*_***u***_*t*_ + ***X***_*t*_ = {*x*_*t*_, *x*_*t*−1_, *u*_*t*_ + *y*_*t*_}, is predicted by a function, ***W***_***simp***_, as follows:

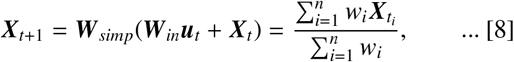

where *n* is the number of nearest neighbors, *t*_*i*_ is the time point of the *i*th nearest neighbor, ***W***_*in*_ is an input weight matrix, and *w*_*i*_ is a weight based on the distance between the target vector to nearest neighbors. The weight is calculated as follows: 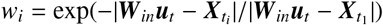 where |·| denotes Euclidean distance. ***W*** _*simp*_, the near future prediction based on behaviors of nearest neighbors in the reconstructed state space, is called “simplex projection” (Sugihara & May 1990), and the numerical method to predict ecosystem’s response based on SSR and simplex projection is called “scenario exploration” (Deyle *et al*. 2013). From an RC point of view, ***W*** _*simp*_ may be applied either before or after the perturbation, that is, either of ***X***_*t*+1_ = ***W***_*in*_***u***_*t*_ + ***W*** _*simp*_ ***X***_*t*_, or ***X***_*t*+1_ = ***W*** _*simp*_(***W***_*in*_***u***_*t*_ + ***X***_*t*_) may work as ERC. The collection of the readout, ***X***_*t*_, was trained using a ridge regression (*λ* = 0.05).

The performance of *in silico* ERC was evaluated using three tasks: Prediction of chaotic dynamics, emulation of nonlinear autoregression moving average (NARMA) time series (Atiya & Parlos 2000), and generation of an autonomous system (Mackey-Glass equation). Detailed information about the parameters and the setting of the numerical experiments is described in Supplementary Text.

### Real-time ERC: A target unicellular microbe

In real-time ERC, real-time ecological dynamics is used as a reservoir. In the present study, the population dynamics of *Tetrahymena thermophila* was used as a reservoir. *Tetrahymena thermophila* (hereafter, *Tetrahymena*) is a unicellular, eukaryotic organism (belongs to ciliates) of which the cell size is *ca*. 30-100 *µ*m. *Tetrahymena* is commonly found in a freshwater ecosystem, and is widely used as a model organism in molecular biology studies. *Tetrahymena* can easily be cultured using a wide variety of media, chambers, and conditions, and its doubling time is *ca*. two hours under an optimal conditions (Cassidy-Hanley 2012).

In reality, an ecological reservoir should have several desired characteristics: (i) reservoir states are easily monitored, and (ii) reservoir states change relatively quickly in response to external forces. As for the first characteristic, *Tetrahymena* population is known to respond to temperature. Its population growth rate maximizes *ca*. 37°C. Our experiment confirmed the response of the *Tetrahymena* population to the medium temperature (Fig. S4). The doubling time is 125 min at 30°C using modified Neff medium (for the medium preparation, see the following paragraph). Also, the *Tetrahymena* population changes its growth rate in response to medium concentration (Fig. S5). These facts indicate that parameters of *Tetrahymena* population dynamics change easily in response to external forces, which may a suitable characteristic as an ecological reservoir. As for the second characteristics, *Tetrahymena* cell size is 30–100 *µ*m, which is relatively large compared with unicellular, prokaryotic organisms. Therefore, their cells can easily be observed under a standard optical inverted microscope. By combining time-lapse imaging and simple image analysis, their cell numbers can easily be monitored (Fig. S6). We chose the *Tetrahymena* population as a candidate for real-time ERC because of these characteristics.

*Tetrahymena thermophila* (strain CU428.2, RRID:TSC_SD00178) was obtained from the *Tetrahymena* Stock Center (Cornell University; https://tetrahymena.vet.cornell.edu/). As described previously (Cassidy-Hanley 2012), *Tetrahymena* is maintained at 27°C in PPYG medium (0.4% Bacto proteose peptone, 0.2% Bacto yeast extract, and 1% glucose). In the PPYG medium, 1% antibioside (antibiotic-antimycotic mixed solution; Nacalai tesque, Kyoto, Japan) was added to prevent growth of harmful fungi. The *Tetrahymena* populations were transferred to fresh medium once per week. When used for experiments, 50 *µ*l of the *Tetrahymena* stock was transferred to modified Neff medium (0.25% Bacto proteose peptone, 2.5% Bacto yeast extract, and 33.3 *µ*M FeCl_3_).

### *Real-time ERC: Experiments using a* Tetrahymena *population*

The computational capability of RC positively correlates with the reservoir size, and to increase the reservoir size, we used three concentrations of modified Neff medium in the experiments. For the 1.6% medium condition (“Low nutrient” in Fig. 3F), 50 *µ*l of the stock *Tetrahymena*, 200 *µ*l of modified Neff, and 4750 *µ*l of H_2_O were mixed and pre-incubated overnight (*ca*. 20 hours) at 27°C. Then, two ml of the pre-incubated medium and three ml of H_2_O were mixed and used for the experiments. For the 4% medium condition (“Med. nutrient” in Fig. 3F), 50 *µ*l of the stock *Tetrahymena*, 200 *µ*l of modified Neff, and 4750 *µ*l of H_2_O were mixed, pre-incubated for three hours at 30°C, and used for the experiments. For the 10% medium condition (“High nutrient” in Fig. 3F), 50 *µ*l of the stock *Tetrahymena*, 500 *µ*l of modified Neff, and 4450 *µ*l of H_2_O were mixed, pre-incubated for three hours at 30°C, and used for the experiments. For each medium condition, two separate runs were performed to confirm the reproducibility and ESP of the population dynamics of *Tetrahymena* (Figs. 3 and S8).

The *Tetrahymena* population in the medium was incubated in an aluminum chamber (Fig. 3A–D), and the temperature inside the aluminum chamber was automatically regulated using a custom temperature regulator system (E5CC; OMRON, Kyoto, Japan). This system enables accurate temperature control of the medium inside the aluminum chamber, and a user can set a maximum of 256 consecutive temperature values at flexible time intervals. In the experiments, medium temperatures were set between *ca*. 10–25°C because the population growth coefficient responds well to the medium temperature in this temperature range (Fig. S4). Medium temperatures were changed every five mins, and thus the total incubation time for each experiment was 256 time-steps × 5 mins = 1280 mins. The medium temperature was also monitored every minute using a temperature logger/sensor (Ondotori TR-52i; T&D, Matsumoto, Japan). We used four time series as inputs, ***u***_*t*_: (i) uniform random, (ii) Lorenz attractor, (iii) empirical fish-catch time series (flatfish; *Paralichthys olivaceus*), and (iv) empirical fish-catch time series (Japanese jack mackerel; *Trachurus japonicus*). The first one was used to quantify the memory capacity of the ecological reservoir, and the other three were used to test the predictive capability of the *Tetrahymena* reservoir.

During the incubation, images of the *Tetrahymena* population at the bottom of the aluminum chamber were taken every minute, resulting in 1280 images for each run. The number of cells in each image was semi-automatically counted using Fiji (Schindelin *et al*. 2012) and OpenCV (Bradski 2000) with custom python codes (Fig. S6). Briefly, the color images were converted to gray scale images and background was subtracted using the rolling-ball algorithm implemented in Fiji. Then, the gray-scale, background-subtracted images were binarized, and the *Tetrahymena* cells in each image were identified using the watershed algorithm.

### Real-time ERC: Preprocessing of cell count data and training of readout

By using the automated temperature regulator and cell counting system, responses of the *Tetrahymena* population to changing medium temperatures were monitored semi-automatically (for an example of the population dynamics, see Fig. S6B). Although there is only a single species in the system, the population dynamics at the bottom of the aluminum chamber is a result of complex interactions among biotic and abiotic factors such as medium temperature, cell physiological states, cell-cell interactions, and behaviors. Indeed, previous studies demonstrated that *Tetrahymena* population dynamics and behavior may be influenced by temperature and medium concentrations, and complex nonlinear interactions seem to govern the dynamics and behavior (Jordan *et al*. 2013; Weisse *et al*. 2016). In addition, another study showed that, using an experimental prey-predator system, *Tetrahymena pyriformis* may exhibit various population dynamics from simple equilibrium to limit cycle to chaotic dynamics (Becks *et al*. 2005). These studies imply that complex, but deterministic, nonlinear interactions drive the population dynamics of *Tetrahymena*, and that such dynamics can be a reservoir that processes information efficiently.

To use the cell count data as reservoir states, we processed the raw data. First, a long-term trend of the cell count data was removed by calculating residuals of a general additive model (Fig. S6B,C). This process was done to make the time series stationary. Second, the residuals were divided by the predicted values by the additive model at the same time point (Ten was added to the predicted values to mitigate the effect of a low cell count) (Fig. S6D). This process was done to correct the dependence of fluctuations in the population dynamics on the absolute cell density. The resulting time series includes information about how the *Tetrahymena* population responds (i.e., an index of the relative response) to the medium temperature and was used as a reservoir state (Fig. S6D).

We obtained two preprocessed time series of the *Tetrahymena* population for each medium concentration for each input time series (Figs. 3B and S8). Therefore, we had six preprocessed time series for each input time series. Each preprocessed *Tetrahymena* time series, ***X***_*t*_, has 1280 values with a one-minute time interval, e.g., ***X***_*t*_ = {*x*_1_, *x*_2_, …, *x*_1280_}. On the other hand, the input value, ***u***_*t*_, has 256 values with a five-minute time interval, ***u***_*t*_ = {*u*_1_, *u*_2_, …, *u*_256_}. In the analysis, we time-multiplexed the reservoir states. For example, five individual reservoir states {*x*_1_, *x*_2_, …, *x*_5_} correspond to one input value, *u*_1_, enabling us to increase the reservoir size.

For explanation purposes, we name the reservoir state 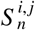, where *i, j*, and *t* indicate a nutrient condition (“*l*,” “*m*,” and “*h*” denote low, medium, and high, respectively), a replicate of the experiment (“1” or “2”), and time step, respectively. For example, 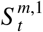 indicates the reservoir state taken from the first run of the medium nutrient concentration (4% modified Neff). For the uniform random value inputs, 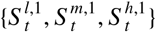 was used for the training and 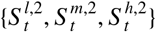 was used for the testing. As each *Tetrahymena* time series, 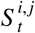, was time-multiplexed for each run, the combined reservoir state, 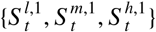, has a 256 × 15 matrix. ***W***_*out*_, a 1 × 15 matrix, was learned by a ridge regression, and used to predict a past input value with the test reservoir state, 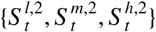. For the prediction tasks, all six reservoir states were time-multiplexed and combined. Thus, 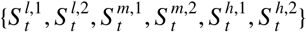 is a 256 × 30 matrix. ***W***_*out*_ was learned by a ridge regression, and the remaining data were used for testing. Detailed information on the size of training and testing data is described in Supplementary Text.

### Computation

Analysis codes to reproduce the results will be available at Github (https://github.com/ong8181/ecological-RC). For RC, custom codes written in python 3.6.10 executed in R environment with “reticulate” (version 1.18) (Ushey *et al*. 2020) were used. Image processing for cell counting was performed using Fiji (version 2.1.0/1.53c) (Schindelin *et al*. 2012) and OpenCV (version 4.4.0) (Bradski 2000) with custom python codes. Preprocessing of the cell count data was performed using “tidyverse” (version 1.3.0) (Wickham 2017), “lubridate” (version 1.7.9.2) (Grolemund & Wickham 2011), and “mgcv” (version 1.8.34) (Wood 2004) packages of R4.0.3 (R Core Team 2020).

## Data and code availability

Empirical ecological time series and analysis scripts will be available at https://github.com/ong8181/ecological-RC. The citable versions of the supplementary videos are available at figshare: Supplementary Video 1 (https://doi.org/10.6084/m9.figshare.16608802) and Supplementary Video 2 (https://doi.org/10.6084/m9.figshare.16608808).

## Acknowledgements

We thank Ai Matsuda for help with the figure editing, and Sayaka Suzuki for help with the incubation experiment.

## Author Contributions

MU conceived the idea; MU and KN designed the research; MU and KW collected the long-term ecological time series from the field; MU, YT, and KN performed the computation for *in silico* ecological reservoir computing (ERC); MU and YF designed the incubation experiment; MU performed the real-time ERC experiment with help from YF; and MU and KN analyzed the data from the real-time ERC experiment. MU and KN wrote the first draft and completed the final manuscript with contributions from all coauthors.

