## Supplementary Figures for "Computational capability of ecological dynamics"

### Contents:

Figure S1. Results of the demonstration of *in silico* Ecological Reservoir Computing using a toy model

Figure S2. *In silico* Ecological Reservoir Computing using scenario exploration

Figure S3. Measurements of information processing capacity of *in silico* ecological reservoirs

Figure S4. Dependence of *Tetrahymena* population growth rate on the medium temperature

Figure S5. Dependence of *Tetrahymena* population growth rate on the medium concentrations

Figure S6. Calculation of reservoir state of real-time Ecological Reservoir Computing

Figure S7. Echo State Property of real-time Ecological Reservoir Computing

Figure S8. Input sequences, medium temperature, and six reservoir states of the *Tetrahymena* ecological reservoir.

Figure S9. Predictions of medium temperature (not input temperature) by real-time Ecological Reservoir Computing

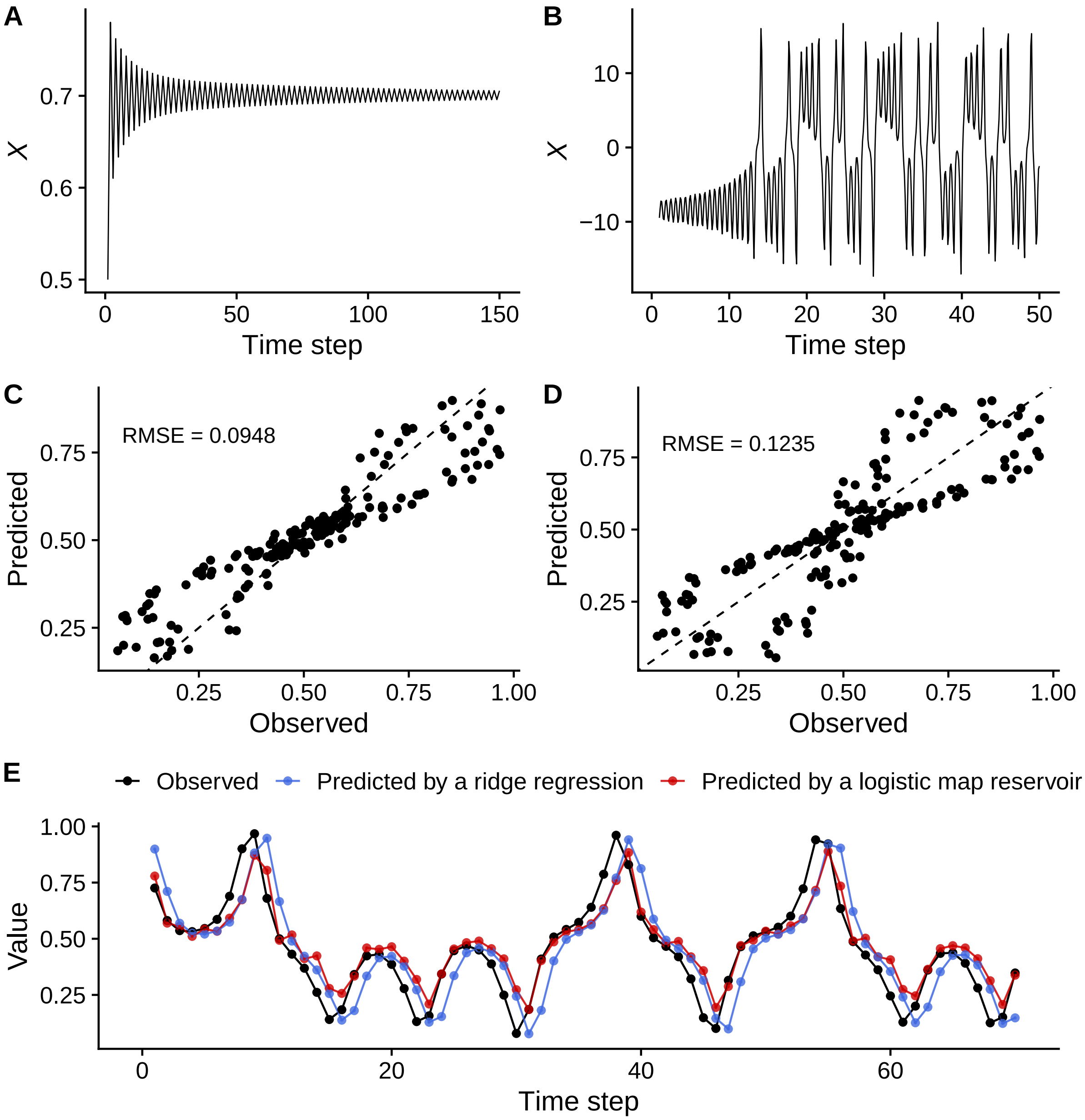


**Figure S1.** (**A**) Time series of a variable *x* of a logistic map. (**B**) Time series of a variable *x* in Lorenz system. The parameter values to generate the dynamics are described in the Supplementary Text (**C**-**E**) Results of reservoir computing using the logistic system as a reservoir. Input vectors are a variable *x* of the Lorenz system, and the one-step prediction accuracy was evaluated by Root Mean Square Error (RMSE). (**C**) The correlation between observed and predicted values by a logistic map reservoir. (**D**) As a comparison, the correlation between observed and predicted values by a ridge regression is shown (i.e., predictions without a reservoir). The prediction accuracy of the logistic reservoir is slightly better than that of the ridge regression. (**E**) Time series plot of the observed and predicted values by the logistic reservoir and ridge regression. The predictions made by a logistic reservoir delayed by the observed values, suggesting that a relatively low prediction capacity of the reservoir due to the small reservoir size ($N = 2$).

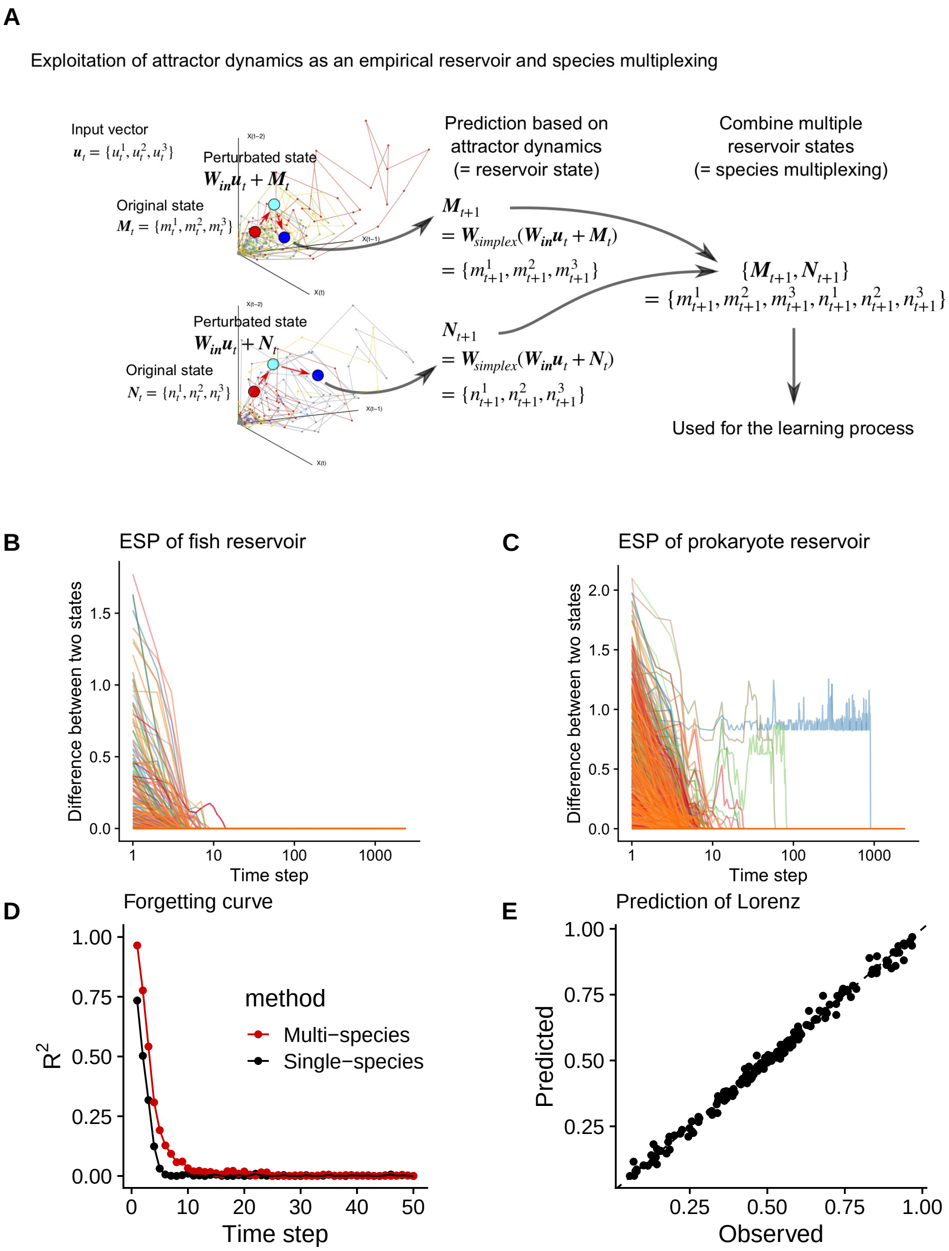


**Figure S2.** (**A**) Schematic illustrations of species-multiplexing using scenario exploration. Simulated ecosystem responses are collected to generate a “multiplexed reservoir state”, which is used for the learning process. Red point in the reconstructed state space indicates the original vector, light blue point indicates perturbated vector, and blue point indicates the predicted response of the system by simplex projection. (**B**) Echo state property (ESP) of the reconstructed fish reservoirs (47 fish species included). For each run, the computation of *in silico* ERC started from two different initial conditions, and the dependence of the state difference on the time step was measured by the Euclidean distance between the two state. For each species, the numerical experiment repeated five times. (**C**) ESP of the reconstructed prokaryote reservoirs (500 prokaryote species included). $y$-axis in **B** and **C** indicates the difference between reservoir states started from different initial conditions and the difference converges to zero when the same input sequence is used. (**D**) An example of forgetting curves of reconstructed prokaryote reservoir. Forgotten curves of single-species prokaryotic reservoir (*Bdellovibrio* sp.) and species-multiplexed prokaryotic reservoir are shown. (**E**) The one-step prediction accuracy of Lorenz system predicted by species-multiplexed reservoir evaluated by the correlation between predicted and observed values.

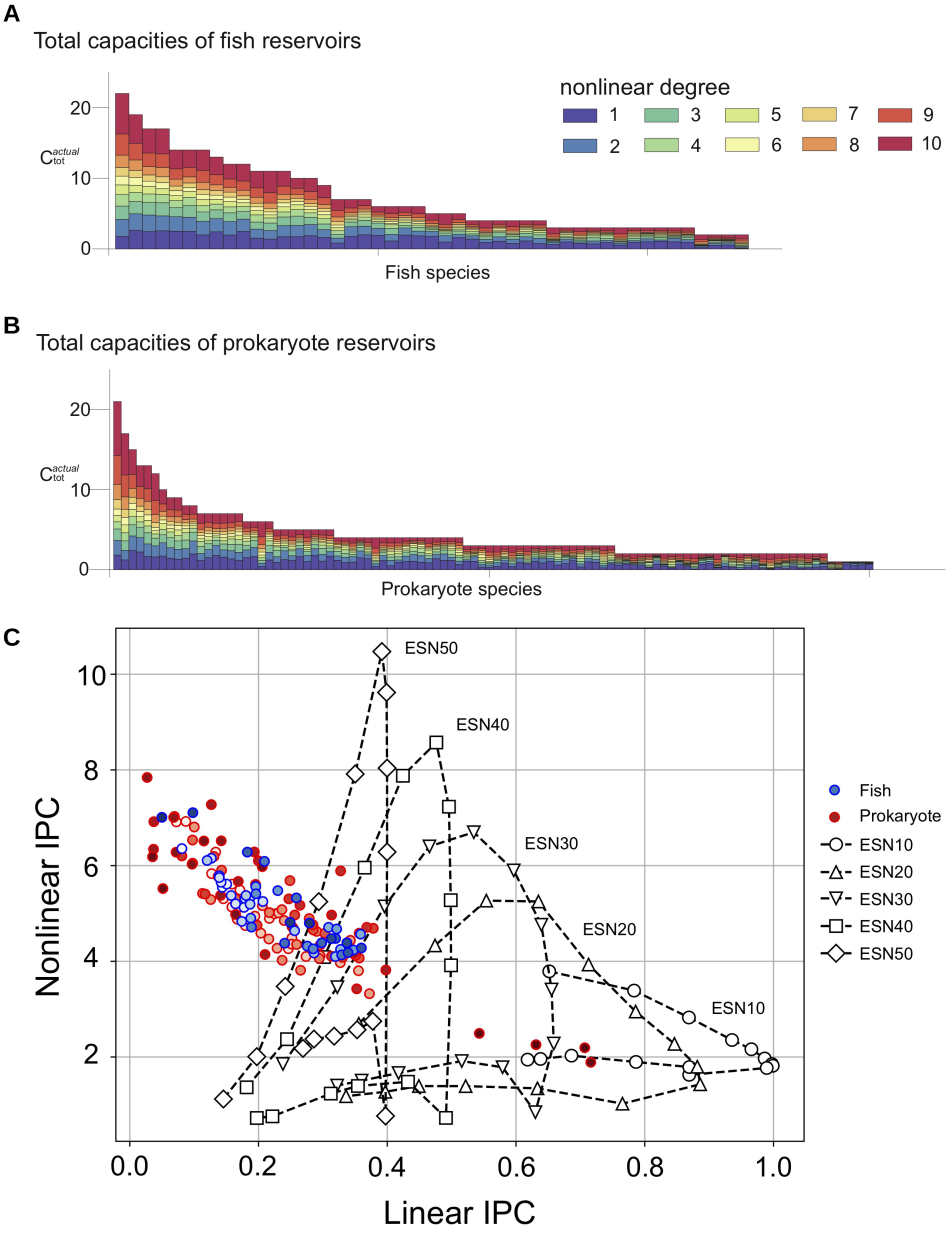


**Figure S3.** Measurements of information processing capacity of *in silico* ecological reservoirs. **A** and **B** show the actual total capacities and the degree of nonlinearity consisting of each capacity for each species in fish reservoirs (**A**; top 47 species are shown) and prokaryote reservoirs (**B**; top 100 species are shown). Horizontal axis shows the sorted number of species in descending order of their capacity size. Colors indicate the degree of nonlinearity. **C** shows the linearity-nonlinearity balance of the actual total IPC. Horizontal axis shows $L(C_{tot}^{actual})$, and vertical axis shows $NL(C_{tot}^{actual})$. Results from all the species were overlaid together with the results of the echo state networks (ESNs) for comparison. Different symbols indicate different reservoir settings (*in silico* ERC or ESN), and different colors of the filled circles indicate different species. For the setting of ESNs, we used a nonlinear activation function ($f(x) = tanh\ x$), the input weight was randomly assigned in the scale of [-0.01, 0.01], and the spectral radius of the internal weight matrix of the ESN was altered from 0.1 to 1.5 for ESNs. The number of nodes for the ESNs are varied from 10 to 50 for comparison, and all the points and lines show the averaged value for 10 trials obtained by randomly choosing the weights of ESNs with respect to the given condition.

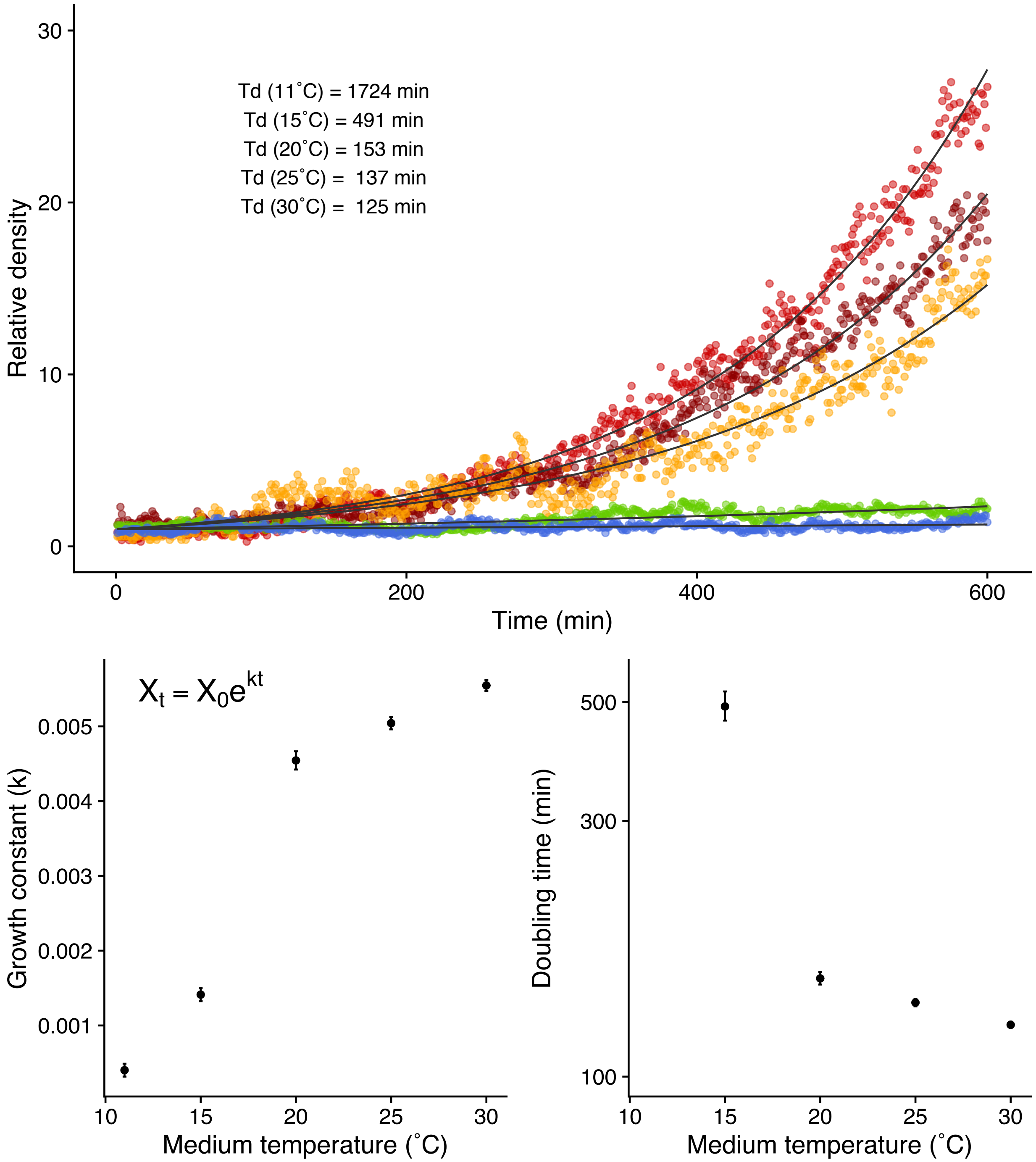


**Figure S4.** Dependence of *Tetrahymena* population growth rate on the medium temperature. (**A**) Changes in population density under different temperature conditions. The initial population density is scaled to one ("Relative density") to facilitate the comparison among different temperatures. Detailed experimental conditions are described in the text. (**B**) Growth constant and medium temperature. Growth constants were estimated by fitting the data to the exponential growth model. (**C**) Doubling time ($T_d$) and medium temperature ($T_d = ln(2)/k$).

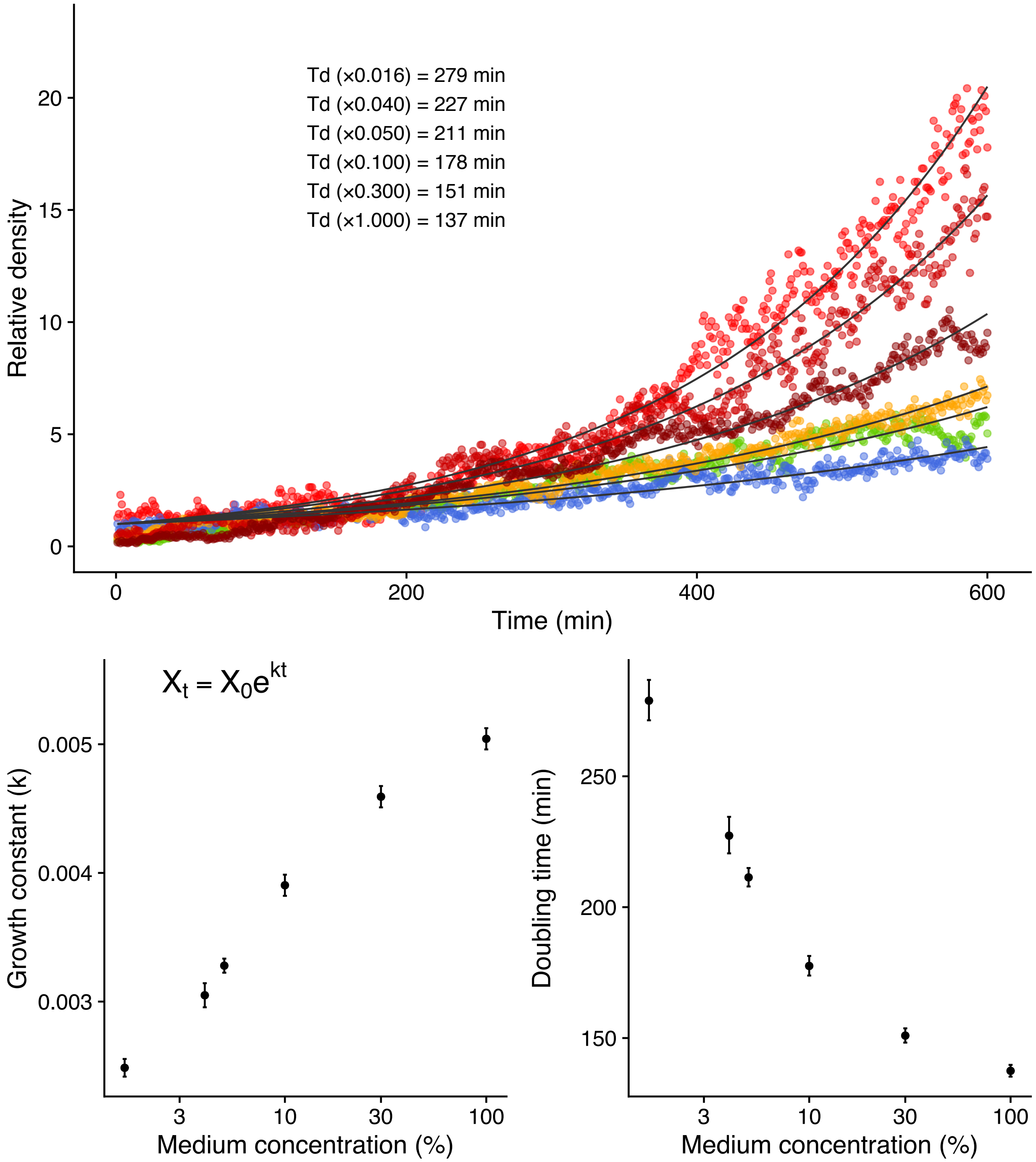


**Figure S5.** Dependence of *Tetrahymena* population growth rate on the medium concentrations. (**A**) Changes in population density under different medium concentrations. The initial population density is scaled to one ("Relative density") to facilitate the comparison among different temperatures. Detailed experimental conditions are described in the text. (**B**) Growth constant and medium concentration. Growth constants were estimated by fitting the data to the exponential growth model. (**C**) Doubling time ($T_d$) and medium concentration ($T_d = ln(2)/k$).

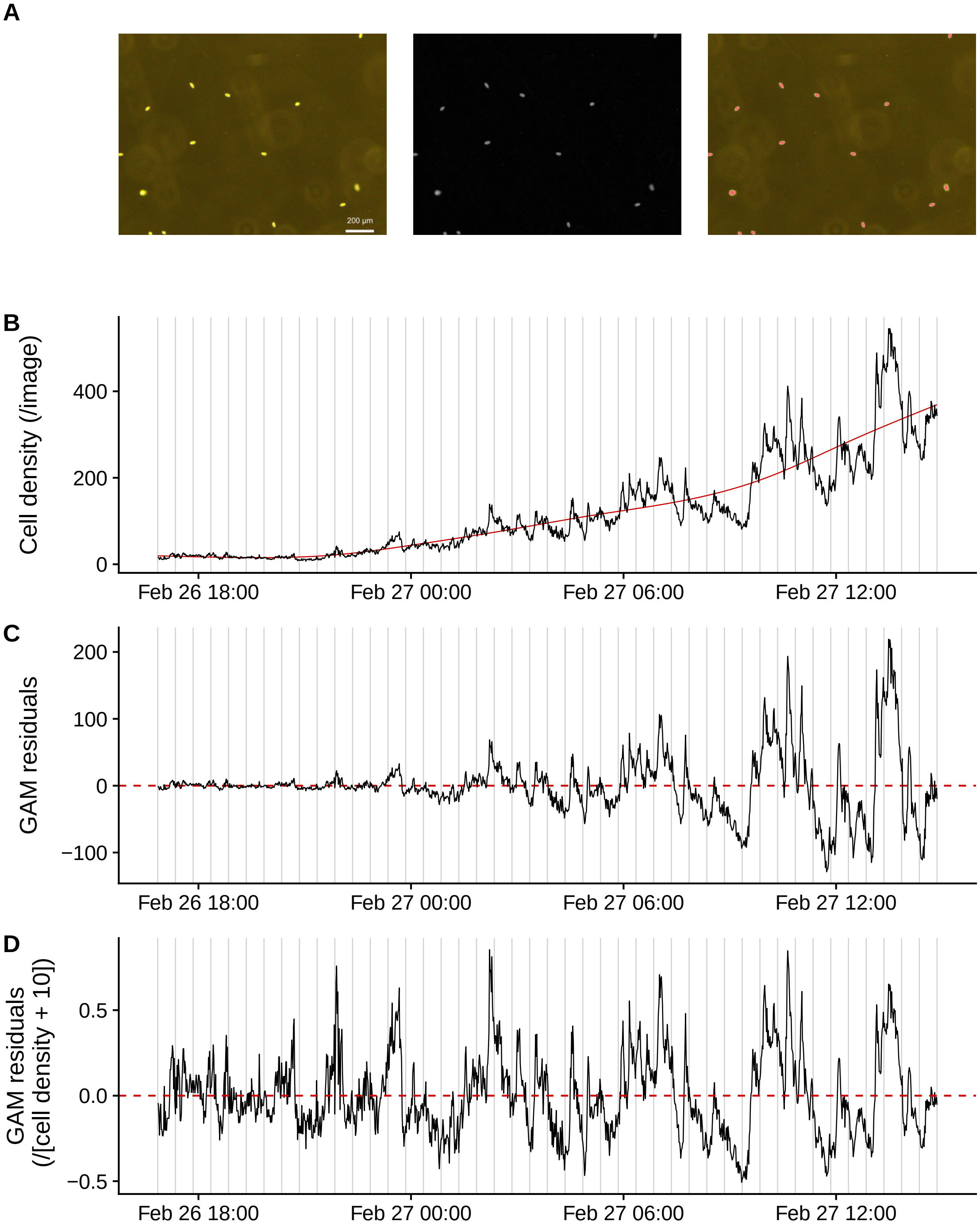


**Figure S6.** Calculation of reservoir state of real-time ecological reservoir computing (ERC). (**A**) Processing of *Tetrahymena* cell images. Images (×40) were obtained using an optical inverted microscope (the image covers *ca*. 1.8 mm width × 1.35 mm height). The background of the image was subtracted, converted to the gray-scale, and cells were counted by the watershed algorithm. (**B**) Change in the *Tetrahymena* cell density. Solid red line indicates a trend estimated by general additive model (GAM regression). (**C**) Residuals of GAM regression. Dashed red line indicates zero. (**D**) Relative GAM residuals. GAM residuals divided by [cell density + 10] to stabilize the fluctuations, which is used as reservoir state of real-time ERC.

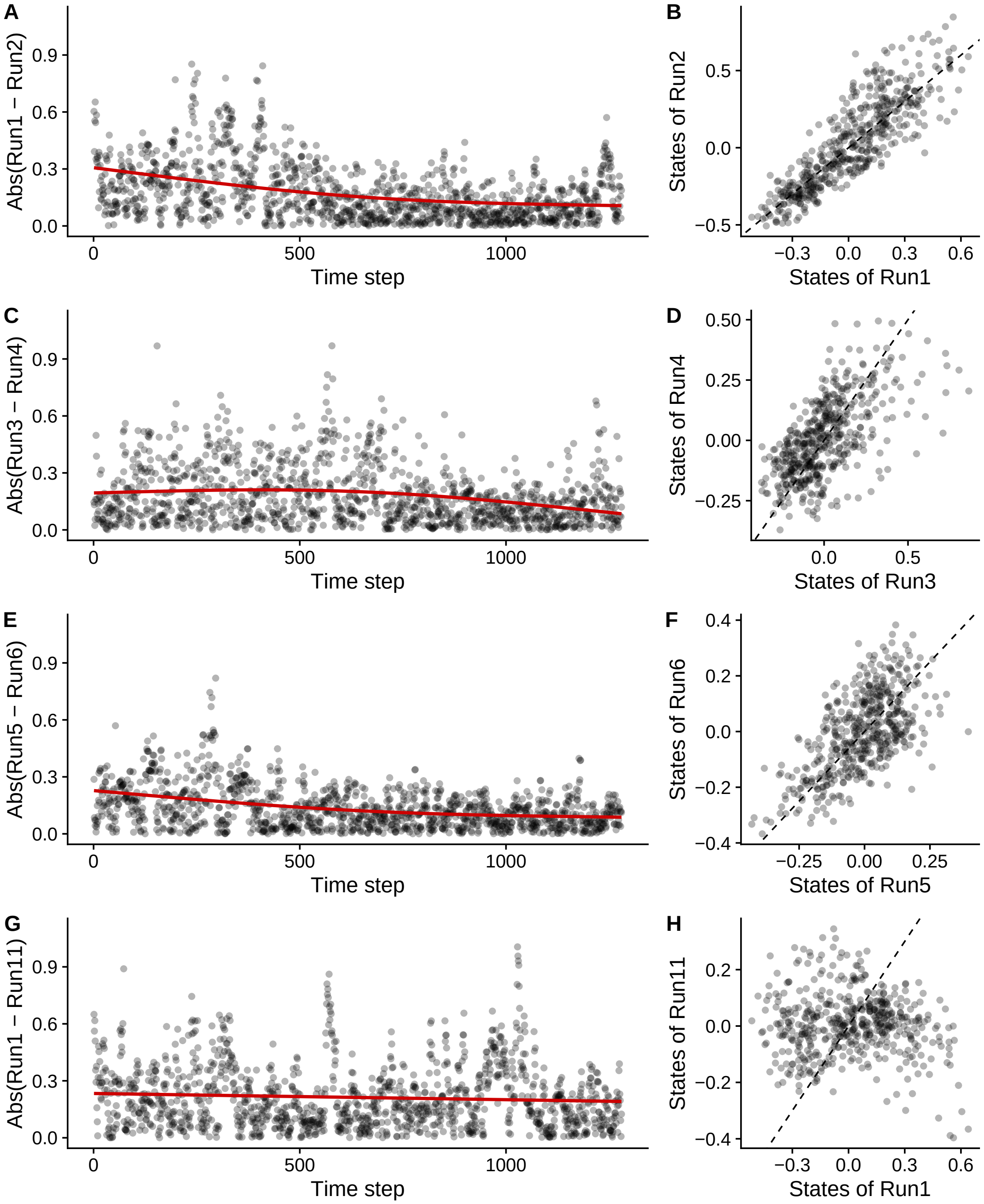


**Figure S7.** Echo state property (ESP) of real-time ecological reservoir computing (ERC). Two runs for each medium concentration were tested and the input sequence was identical for (**A**–**F**). (**A**, **B**) Comparison of two reservoir states (i.e., relative GAM residuals explained in Fig. S5; Run1 and Run2) where the same input sequence was added in 4% Neff medium. Time series plot (**A**) and scattered plot (**B**). (**C**, **D**) Comparison of two reservoir states in 1.6% Neff medium (Run3 and Run4) and (**E**, **F**) in 10% Neff medium (Run5 and Run6). (**G**, **H**) Comparison of two reservoir states in 4% Neff medium, but the input sequence was different for Run1 and Run11. Reservoir outputs converged for the identical inputs when the medium concentration was the same (**A**-**F**). On the other hand, the reservoir outputs did not converge when the input sequences were different (**G**, **H**). Red and dashed lines indicate GAM regression and 1:1 line, respectively.

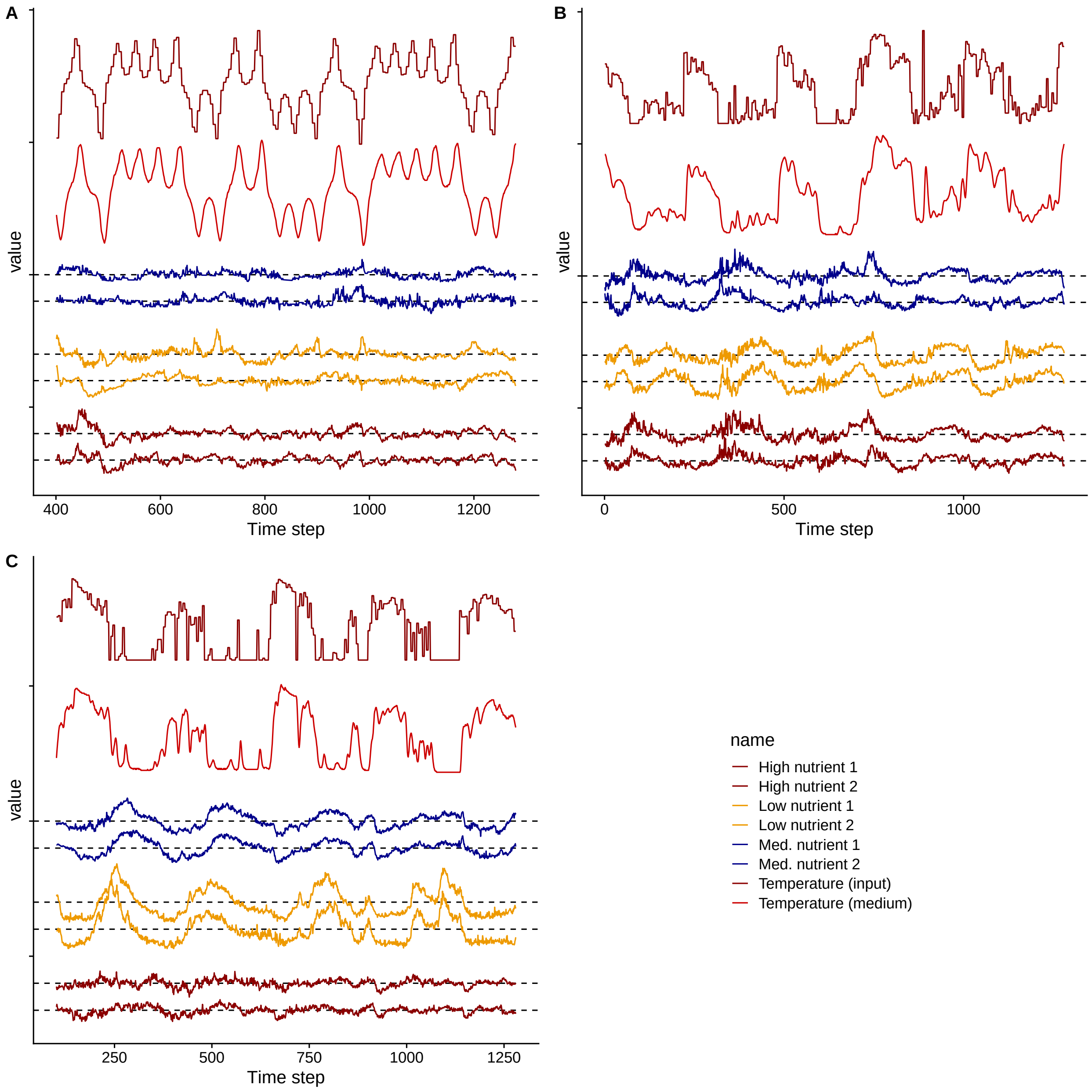


**Figure S8.** Input sequences, medium temperature, and six reservoir states of ecological reservoir. (**A**) Input sequence that represents Lorenz system and the responses of *Tetrahymena* reservoir. (**B**) Input sequence that represents *Paralichthys olivaceus* time series and the reservoir responses. (**C**) Input sequence that represents *Trachurus japonicus* time series and the reservoir responses. The first top and second top time series indicate input temperature and medium temperature, respectively. The dark blue (third and fourth top), orange, and dark red lines indicate reservoir states of middle (4%), low (1.6%) and high (10%) medium conditions, respectively.

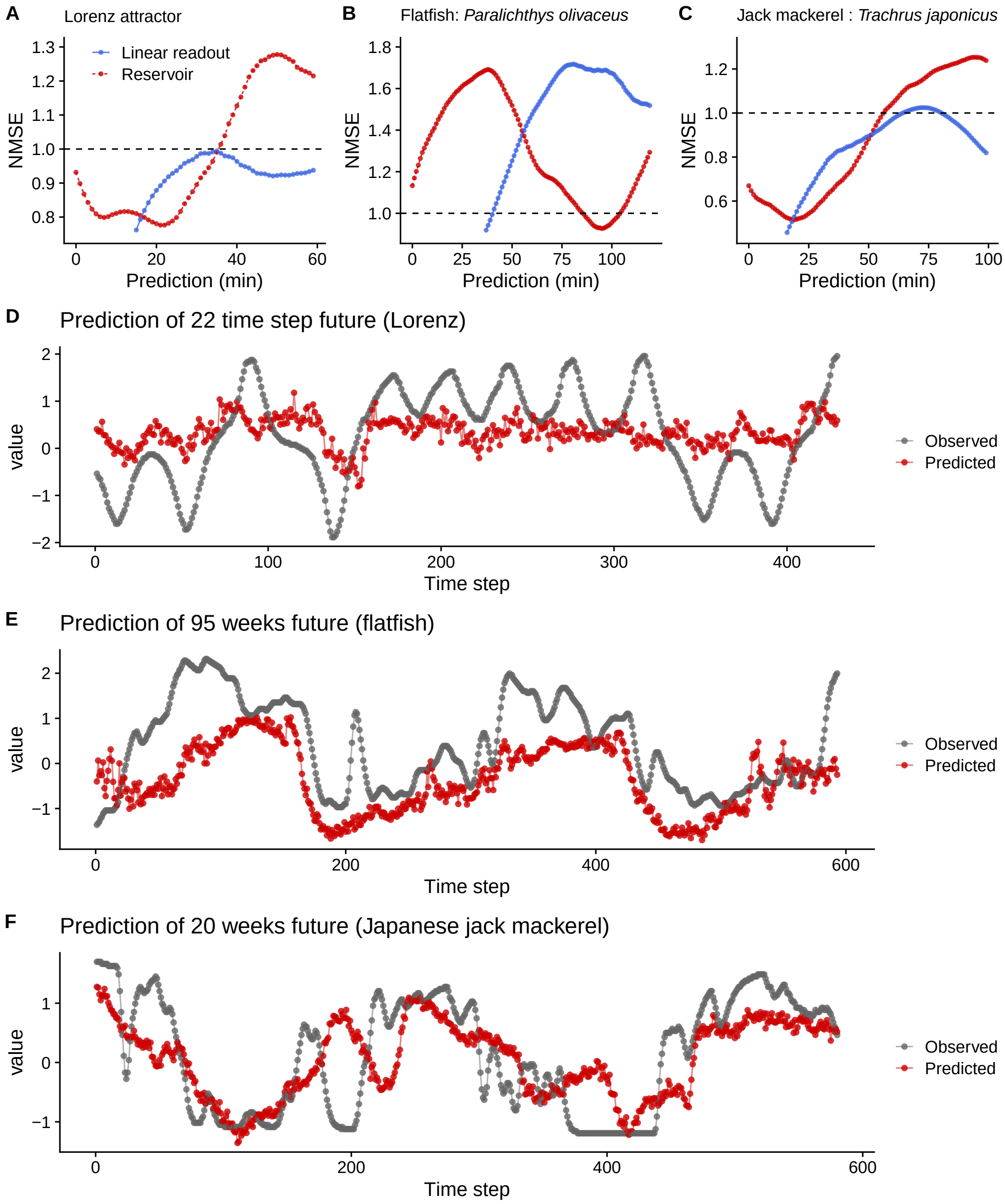


**Figure S9.** Predictions of medium temperature (not input temperature) by real-time ecological reservoir computing (ERC), the *Tetrahynema* reservoir. Instead of using values set to the temperature regulator, medium temperatures were used as "input" sequences. (**A**, **B**, **C**) Prediction accuracy (Normalized Mean Square Error; NMSE) of Lorenz system (**A**), *Paralichthys olivaceus* time series (**B**), and *Trachurus japonicus* time series (**C**) by the real-time ERC. Red points and line indicate predictions made by the real-time ERC, and blue points and line indicate temporal autocorrelation. $x$-axis indicates prediction time horizon. For all cases, predictions made by ERC are better than a simple ridge regression for at least some prediction time steps, suggesting that ERC works as an effective reservoir system. (**D**, **E**, **F**) Time series of the predictions. Gray points and line indicate true values (observed values), and red points and line indicate predictions by ERC.
