## Supplementary Text for "Computational capability of ecological dynamics"

### Contents:

1. Detailed information and parameterizations of *in silico* Ecological Reservoir Computing
2. Measurements of information processing capacity of *in silico* ecological reservoirs
3. Preparation of media for *Tetrahymena thermophila*
4. Measurements of growth parameters of *Tetrahymena* population
5. Setting of ERC using *Tetrahymena* population dynamics

### 1. Detailed information and parameterizations of *in silico* Ecological Reservoir Computing

#### *Empirical ecological time series*

Two types of empirical ecological time series were used for *in silico* ecological reservoir computing (ERC): Count-based, fish catch time series taken from pelagic regions in Japan, and DNA-based, prokaryotic time series taken from artificial rice plots.

Count-based fish catch time series were collected mainly by checking a report of fish catch on a fishing boat which was announced on a website. Briefly, approximately 800–1500 fishing boat websites reported their daily fish catch, fish size, and fish species, and most of the information has been manually collected by staff at B.Creation Inc. every day. In addition, some fishing boat owners may report their daily fish catch information to B.Creation Inc. via an online or manual data submission system. In total, the fish catch information from *ca.* 2500 fishing boats has been continuously monitored. The collected count data were converted to fish catch per unit effort and compiled as weekly time series, which were used for *in silico* ERC.

DNA-based, prokaryotic time series were described in detail in a previous study (Ushio 2020). Briefly, five 90 cm × 90 cm small rice plots were established in 2017 in an experimental field in Center for Ecological Research, Kyoto University. Water samples were taken daily from the rice plots from May to September 2017 (122 days), and 610 water samples were collected in total (122 days × 5 plots). Prokaryotic DNAs in the water samples were quantitatively measured using a quantitative DNA metabarcoding approach (Ushio 2019). Among the prokaryotic species detected, 500 abundant and frequently detected species were selected and further analyzed in the previous study (Ushio 2020). Time series of these 500 prokaryotic species were also used in this study.

#### *Lorenz attractor prediction by species-multiplexed empirical ecological reservoir*

Lorenz system is described as follows:

$$\frac{dx}{dt} = -px + py$$
$$\frac{dy}{dt} = -xz + rx - y$$
$$\frac{dz}{dt} = xy - bz, \qquad \text{... [9]}$$

where $(p, r, b)$ were set to $(10, 28, 8/3)$. The differential equations were solved using “deSolve” package (Soetaert *et al.* 2010) of R. Initial values of $x$, $y$, and $z$ were set to one and time steps were set from 0 to 100 with 0.01 intervals.

For species-multiplexed *in silico* ERC, time series of the 500 prokaryotic species were used as ecological reservoirs. Optimal embedding dimensions ($E$) of the prokaryotic time series were estimated by simplex projection, and the total reservoir size ($N$) was 3271 when the 500 species were multiplexed (i.e., the sum of $E$ of each species corresponds to total reservoir size). The sparsity (= the proportion of zero in the matrix elements) of $\boldsymbol{W}_{in}$, $\boldsymbol{W}_{sparsity} = 0$. Matrix elements of $\boldsymbol{W}_{in}$ were multiplied by two to adjust the influence of input vectors ($\boldsymbol{W}_{strength} = 2$). Readout was trained using a ridge regression ($\lambda$ for the regularization = 1). The optimal values for $\boldsymbol{W}_{in}$ and $\boldsymbol{W}_{sparsity}$ were determined by a grid search.

#### *NARMA emulation by species-multiplexed empirical ecological reservoir*

Nonlinear autoregression moving average (NARMA) time series were used to evaluate the emulation capacity of *in silico* ERC (Atiya & Parlos 2000). The first NARMA system is the following second-order nonlinear dynamical system:

$$y_{t+1} = 0.4\,y_t + 0.4\,y_t\,y_{t-1} + 0.6\,I_t^3 + 0.1, \qquad \text{... [10]}$$

where $y_t$ denotes the output of the system. For descriptive purposes, we call this system NARMA2. The second NARMA system is the following nonlinear dynamical system that has an order of $n$:

$$y_{t+1} = \alpha\,y_t + \beta\,y_t \sum_{j=0}^{n-1} y_{t-j} + \gamma\,I_{t-n+1}\,I_t + \delta, \qquad \text{... [11]}$$

where $t$ denotes time, and $(\alpha, \beta, \gamma, \delta)$ are set to $(0.3, 0.05, 1.5, 0.1)$. Here, $n$ is varied for values of 3, 4, 5, and 10 and the corresponding systems are called NARMA3, NARMA4, NARMA5, and NARMA10, respectively. The input $I_t$ is expressed as a product of three sinusoidal functions with different frequencies:

$$I_t = 0.2\sin(2\pi f_1 \frac{t}{T})\,\sin(2\pi f_2 \frac{t}{T})\,\sin(2\pi f_3 \frac{t}{T}), \qquad \text{... [12]}$$

where $t$ denotes time, and $(f_1, f_2, f_3, T)$ is set to $(2.11, 3.73, 4.33, 400)$, and the parameter $T$ controls the phase velocity of the input time series. For all NARMA time series, the initial value is set to 0.2. For the emulation task, 7000 time points were generated, and the first 6000 time points were discarded, and the remaining 1000 time points were used. Among the 1000 time points, 500 time points were used for the training, and the remaining 500 points were used for testing.

For species-multiplexed *in silico* ERC, the reservoir dynamics is $\boldsymbol{X}_{t+1} = \boldsymbol{W}_{simp}(\boldsymbol{W}_{in}u_t + \boldsymbol{X}_t)$, namely, $\boldsymbol{W}_{simp}$ was applied after perturbations were added. Following parameter values were used: the total reservoir size ($N$) = 3271, $\boldsymbol{W}_{sparsity}$ = 0, and $\boldsymbol{W}_{strength}$ = 0.5. The optimal values of $\boldsymbol{W}_{sparsity}$ and $\boldsymbol{W}_{strength}$ were determined by a grid search. Readout was trained using a ridge regression ($\lambda$ for the regularization = 1).

Echo state network (ESN) was also used to emulate NARMA system for comparison. The following parameter values were used for ESN: the total reservoir size ($N$) = 3271, $\boldsymbol{W}_{sparsity}$ = 0.6, and $\boldsymbol{W}_{strength}$ = 0.75. $\boldsymbol{W}$ is a random weight matrix for the reservoir dynamics (see Eqn. [6]), the sparsity of $\boldsymbol{W}$ is set to 0.4. Spectral radius of $\boldsymbol{W}$ is set to 0.99. The optimal values of $\boldsymbol{W}_{sparsity}$, $\boldsymbol{W}_{strength}$, and the sparsity and spectral radius of $\boldsymbol{W}$ were determined by a grid search. Readout was trained using a ridge regression ($\lambda$ for the regularization = 1).

#### *Generation of Mackey-Glass attractor by species-multiplexed empirical ecological reservoir*

The Mackey-Glass equation is as follows:

$$y_{t+1} = y_t + \delta\left(\frac{0.2\,y_{t-\tau/\delta}}{(1 + y_{t-\tau/\delta})^{10}} - y_t\right), \qquad \text{... [13]}$$

where $t$ denotes time and $(\delta, \tau)$ is set to $(0.1, 17)$. The initial value is set to 0.5. For the Mackey-Glass task, 7000 time points were generated, and the first 5000 time points were discarded, and the remaining 2000 time points were used. Among the 2000 time points, 1600 time points were used for the training the attractor dynamics, and the remaining 400 points were used for the closed-loop generation of the attractor dynamics.

As for the generation of the Mackey-Glass equation dynamics, the time evolution of the neuronal states in the reservoir is described as:

$$\boldsymbol{X}_{t+1} = (1 - a\,C)\,\boldsymbol{X}_t + C\Big((\boldsymbol{W}_{in}\,\boldsymbol{u}_{t+1} + \boldsymbol{X}_t) + \boldsymbol{X}_{back}\,y_t\Big), \qquad \text{... [14]}$$

where $\boldsymbol{u}_t$ is a constant input and set to 0.2 and $(a, C)$ is set to $(0.44, 0.9)$. $\boldsymbol{W}_{back}$ is a uniform random matrix that represents feedback link from the output to the reservoir.

For species-multiplexed *in silico* ERC, the following parameter values were used: the total reservoir size $(N) = 3271$, $\boldsymbol{W}_{sparsity} = 0$, and $\boldsymbol{W}_{strength} = 0.1$. Because the Mackey-Glass task is a closed-loop system, the learning process includes feedback link (characterized by a matrix $\boldsymbol{W}_{back}$). The sparsity of $\boldsymbol{W}_{back}$ is set to zero, and matrix elements of $\boldsymbol{W}_{back}$ were multiplied by one to adjust the influence of feedback vectors. The optimal values of $\boldsymbol{W}_{sparsity}$, $\boldsymbol{W}_{strength}$, and the sparsity of $\boldsymbol{W}_{back}$ were determined by a grid search. Readout was trained using a ridge regression ($\lambda$ for the regularization = 0.05).

### 2. Measurements of information processing capacity of *in silico* ecological reservoirs

To evaluate the expressiveness of the ecological reservoirs, we used a measure known as information processing capacity (IPC), which quantifies the type and amount of input history held in the system (Dambre *et al.* 2012). A reservoir is a typical input-driven dynamical system, and it processes input information through its state transition. Let us consider the input $\boldsymbol{u}_t$ to be an i.i.d. and the state of the reservoir to be expressed as $x_t \in \mathbb{R}^N$. If the state can be expressed as a function of only input history $\{\boldsymbol{u}_{t-1}, ..., \boldsymbol{u}_0\}$ (this condition is called echo state property, and this function is called an input echo function Jaeger 2001), we can express the function with a weighted sum of orthogonal polynomials of input history (Kubota *et al.* 2019). Here, let $z_{i,t}$ be the $i$th multivariate orthogonal polynomial; then the IPC is defined as the capability of the reservoir state to approximate the polynomial in terms of the normalized mean square error of predicted output as follows:

$$C_i = 1 - \frac{\min\langle (z_{i,t} - \omega^T x_t)^2 \rangle}{\langle z_{i,t}^2 \rangle}, \qquad \text{... [15]}$$

Where $C_i$ is the IPC for the $i$th polynomial, while $\langle\cdot\rangle$ and $\omega \in \mathbb{R}^N$ denote the time average and weight vector of linear regression, respectively. The total capacity $C_{tot}$, which sums up enough types of capacity, is defined as follows:

$$C_{tot} = \sum_{i=1}^{\infty} C_i. \qquad \text{... [16]}$$

Given that the state is expressed as an input echo function, the total capacity is equivalent to the number of linearly independent components within the reservoir states, which is equal to the rank of the reservoir state matrix $r$ ($\leq N$), denoted as follows (Dambre *et al.* 2012; Kubota *et al.* 2019):

$$C_{tot} = r. \qquad \text{... [17]}$$

If the state is a function of other variables except for input history (e.g., its original initial state, other dynamical noise, and measurement noise), $C_{tot} < r$ because the polynomials no longer form a complete orthogonal system for this state.

For the actual analysis of ecological reservoirs, we used uniformly distributed random real values in the range [–1, 1] as an input series $\boldsymbol{u}_t$, which consists of $10^6$ timesteps in total ($10^4$ timesteps are used for washout). In total, the top 47 fish species and top 100 prokaryote species in terms of the total abundance were investigated. The normalized Legendre polynomial, a type of orthogonal polynomial, was used corresponding to the uniform distribution of the input series. We tested all combinations of Legendre polynomials $z_{i,t}$, with the maximum delay $\tau_{max}$ for each degree of nonlinearlity $n$ as $\{(n, \tau_{max})\} = \{(1, 20), (2, 20), (3, 10), (4, 10), (5, 10), (6, 10), (7, 10), (8, 10), (9, 10), (10, 10)\}$, expressed as:

$$z_{i,t} = \prod_{\tau=0}^{\tau_{max}} P_{n_i(\tau)}(\boldsymbol{u}_{t-\tau}), \qquad \text{... [18]}$$

where $P_{n_i(\tau)}$ denotes the normalized Legendre polynomial of degree $n_i(\tau)$ for $\boldsymbol{u}_{t-\tau}$ input. The total number of polynomial combinations is denoted as $M$ (Note: We do not consider the combination where the degree of polynomial $n_i(\tau)$ along the delay is zero).

For each species, we measured the IPCs for all combinations of polynomials in the above-mentioned range, sorted them in decreasing order (renumbering each polynomial from one to $M$), and obtained their sum until the total value equaled the rank (until $M_{actual}$ ($\leq M$)) for each ecological reservoir state. We denote the total capacity summed up in this way as $C_{tot}^{actual}$, where $C_{tot}^{actual} (= \sum_{i=1}^{M_{actual}} C_i) \leq r < \sum_{i=1}^{M_{actual}+1} C_i$. The total IPC will be equal to the rank due to ESP of the ecological reservoirs (see Fig. S2B, C and Eqn. [17]).

Figure S3A and B show how the actual total IPC, $C_{tot}^{actual}$, is decomposed into nonlinear capacities for each species in fish and prokaryote reservoirs. The degree of nonlinearlty is calculated as $\sum_{\tau=0}^{\tau_{max}} n_i(\tau)$ for each polynomial and subsequently classified. We can clearly see that each species has a highly nonlinear IPC profile.

To see how the linear and nonlinear IPCs are balanced in the total IPC for each species, we calculated the linearity $L(C_{toto}^{actual})$ and nonlinearity $NL(C_{toto}^{actual})$ for each IPC as follows (Dambre *et al.* 2012):

$$L(C_{tot}^{actual}) = \sum_{i=1}^{M_{actual}} \delta\left(\sum_{\tau=0}^{\tau_{max}} n_i(\tau) - 1\right)\frac{C_i}{r}, \qquad \text{... [19]}$$

$$NL(C_{tot}^{actual}) = \sum_{i=1}^{M_{actual}} \delta\left(\sum_{\tau=0}^{\tau_{max}} n_i(\tau)\right)\frac{C_i}{r}, \qquad \text{... [20]}$$

where the delta function equals to one only when its argument is zero and is otherwise zero. Figure S3C shows the plot for the linearity-nonlinearity balance of the actual total IPC for all the species. We can clearly see that the reservoirs prepared from ecological data show high nonlinearity balance compared to the conventional echo state networks. The capacities for fish and prokaryote reservoirs overlap in the linearity-nonlinearity plot. One possible explanation of the overlap is that the algorithm drives the pattern: the scheme of *in silico* ERC based on the scenario exploration contributes to the linearity-nonlinearity balance. However, all species that show high linearity and low nonlinearity (i.e., the lower right region of the plot) belong to prokaryote, suggesting that species identity might also influence the pattern. How the algorithm and species identity (ecology and life history of the species) contribute to the information processing capacity is an interesting future question.

### 3. Preparation of media for *Tetrahymena*

In the laboratory, *Tetrahymena* populations are maintained at 27°C in PPYG medium. PPYG medium was prepared as follows. We added two g of Bacto proteose peptone, one g of Bacto yeast extract, and five g of glucose to a flask and brought the volume to 500 ml with distilled water. The solution was then autoclaved at 121°C for 20 min. For storage purposes, one ml of antibioside (antibiotic-antimycotic mixed stock solution [× 100]; Nacalai tesque, Kyoto, Japan) was added to 100 ml of PPYG (1% antibioside) to prevent growth of harmful fungi. Approximately 30 $\mu$l of *Tetrahymena* stock solution were transferred to one ml of a fresh PPYG meium with 1% antibioside in a five-ml sterilized tube once per week.

When used for the experiment, 50 $\mu$l of the *Tetrahymena* stock was transferred to modified Neff medium. Modified Neff medium was prepared as follows. We added 1.25 g of Bacto proteose peptone, 1.25 g of Bacto yeast extract, and 2.5 g of glucose to a flask, and brought to a volume of 500 ml with distilled water. Then, 71.15 $\mu$l of 234 mM $FeCl_3$ (originally prepared using 38% $FeCl_3$) was added so that the final Fe concentration was 33.3 $\mu$M, and the solution was autoclaved at 121°C for 20 min.

### 4. Measurements of growth parameters of *Tetrahymena* population

To quantify the response of *Tetrahymena* population to external forces (i.e., temperature and medium concentrations), the growth rates of *Tetrahymena* population were measured under several temperatures and medium concentrations. For experiments with different temperatures, 50 $\mu$l of *Tetrahymena* stock solution was added to 4950 $\mu$l of 100% modified Neff medium. Then, constant medium temperature was set using the temperature regulation system. We quantified growth rates under five different temperatures, 11°C, 15°C, 20°C, 25°C, and 30°C. For experiments with different medium concentrations, 50 $\mu$l of *Tetrahymena* stock solution was added to 4950 $\mu$l of concentration-adjusted Neff medium. The concentrations of modified Neff medium were 1.6%, 4.0%, 5.0%, 10%, 30% and 100%. The medium temperature was set to 25°C in this experiment. For all experiments, monitoring had been continued for at least 22 hours. The population dynamics were measured using exactly the same system and procedures as those used for ERC. The growth constant and doubling time were estimated by fitting the population density to the equation $x_t = x_0\, e^{kt}$, where $x$, $x_0$, $k$, and $t$ represent population density, initial density, growth constant, and time, respectively. Doubling time was calculated as follows: $T_d = \ln(2)/k$.

### 5. Setting of ERC using *Tetrahymena* population dynamics

As described in the Methods, time-multiplexed reservoir states (*Tetrahymena* density index; Fig. S6) observed in different medium concentrations were used to solve tasks. For memory capacity measurement (uniform random inputs), $\{S_t^{l,1}, S_t^{m,1}, S_t^{h,1}\}$, a $256 \times 15$ matrix, was used for training and $\{S_t^{l,2}, S_t^{m,2}, S_t^{h,2}\}$ was used for testing. The first 120 time points were discarded as a transient phase, and the remaining 136 time points were used for ERC. Readout was trained using a ridge regression ($\lambda$ for the regularization = 1). The coefficients of determination ($R^2$) between the uniform random inputs and predicted values were calculated to measure how well ERC recovers the uniform random inputs.

For the prediction tasks (Lorenz system and two empirical time series), $\{S_t^{l,1}, S_t^{l,2}, S_t^{m,1}, S_t^{m,2}, S_t^{h,1}, S_t^{h,2}\}$, a $256 \times 30$ matrix, was used as reservoir states. For Lorenz system, the first 50 time points were discarded as a transient phase, and the remaining 206 time points were used for ERC. The first half of the 206 time points was used for the training, and the remining half of the data was used for the testing. *lambda* for the ridge regression was 0.5. For *Paralichthys olivaceus* time series, the first half of the time series was used for the training, and the remaining half of the time series was used for the testing. $\lambda$ for the ridge regression was one. For *Trachurus japonicus* time series, the first 10% of the time series was used for the training, and the remaining 90% of the time series was used for the testing. $\lambda$ for the ridge regression was one.
